# One-step generation of auxin-inducible degron cells with high-efficiency homozygous tagging

**DOI:** 10.1101/2023.03.26.534268

**Authors:** Shiqian Li, Yafei Wang, Miesje van der Stoel, Xin Zhou, Shrinidhi Madhusudan, Kristiina Kanerva, Van Dien Nguyen, Nazli Eskici, Vesa M Olkkonen, You Zhou, Taneli Raivio, Elina Ikonen

## Abstract

Auxin-inducible degron (AID) technology is powerful for chemogenetic control of proteolysis. However, generation of human cell lines to deplete endogenous proteins with AID remains challenging. Typically, homozygous degron-tagging efficiency is low and overexpression of an auxin receptor requires additional engineering steps. Here, we establish a one-step genome editing procedure with high-efficiency homozygous tagging and auxin receptor expression. We demonstrate its application in 5 human cell lines, including embryonic stem (ES) cells. The method allowed isolation of AID single-cell clones in 10 days for 11 target proteins with >80% average homozygous degron-tagging efficiency in A431 cells, and >50% efficiency for 5 targets in H9 ES cells. The tagged endogenous proteins were inducibly degraded in all cell lines, including ES cells and ES-cell derived neurons, with robust expected functional readouts. This method facilitates the application of AID for studying endogenous protein functions in human cells, especially in stem cells.

## Introduction

Auxin-inducible degradation (AID) is a technology for chemogenetic control of proteolysis.^1^ To apply AID, a destabilizing peptide, or ‘degron’, is tagged to the target protein by genetic engineering. An auxin receptor (such as *Os*TIR1) is expressed in the same cells, functioning as the substrate recognition subunit of Skp1-Cullin1-TIR1 (SCF^TIR1^) ubiquitin ligase complex. Auxin (such as Indole-3-acetic acid, IAA) as a chemical glue bridges the SCF^TIR1^ ubiquitin ligase and the degron-tagged protein, leading to rapid polyubiquitination and proteasomal degradation of the degron-tagged protein.^1,2^ AID shows rapid and efficient targeted protein degradation, avoids secondary and side effects observed during long-term silencing or CRISPR-knockout, and has provided important mechanistic insights into the functions of diverse target proteins.^3–7^ However, severe pitfalls have limited our ability to harness the full potential of AID.

The first challenge is to engineer optimized AID components. The initial AID systems in mammalian cells using the auxin receptor *Os*TIR1 resulted in severe degradation of target proteins before induction, known as basal or leaky degradation.^8–10^ Several strategies have recently been developed to solve this issue. We identified that substitution of *Os*TIR1 by its homolog *AtAFB2* largely avoided basal degradation.^8^ Moreover, *Os*TIR1(F74G) mutant that exploits a bump-and-hole approach developed in plant, showed no leaky degradation.^9,11^ Other AID components have also been tailored, to enable efficient inducible degradation with low inducer concentration. Substituting the degron miniAID by miniIAA7 improved the inefficient inducible degradation with *At*AFB2, and changing the inducer IAA to 5-PH-IAA required a 670-times lower ligand concentration with *Os*TIR1(F74G).^8,9^ Similarly to 5-PH-IAA, cvxIAA and pico-cvxIAA were used at much lower ligand concentrations than IAA with *Os*TIR1(F74G) and *Os*TIR1(F74A) mutants.^12,13^ These recent developments have resolved apparent technical pitfalls of AID components.

The second challenge is to effectively generate AID cells. Engineering cells with AID to deplete endogenous proteins requires two genetic modifications, i.e. homozygous degron-tagging and auxin receptor expression. With the ease of CRISPR-Cas9 for genetic engineering, applications of AID in mammalian cells have become feasible.^8,10^ However, homozygous degron-tagging through CRISPR/Cas9-mediated homology directed repair (HDR) suffers from low efficiency in mammalian cells.^14^ Hence, FACS sorting or drug selection is first used to enrich homozygously tagged cells before single-cell cloning and an additional engineering step is required to introduce the auxin receptor **(Fig. 1a)**.^8^,^10^ Such multi-step procedures are labor-intensive, time consuming, and cannot be easily scaled up. Moreover, homozygous tagging in pluripotent cells, such as human embryonic stem (hES) cells, suffers from even lower efficiency with limited success.^15,16^

**Figure 1:**
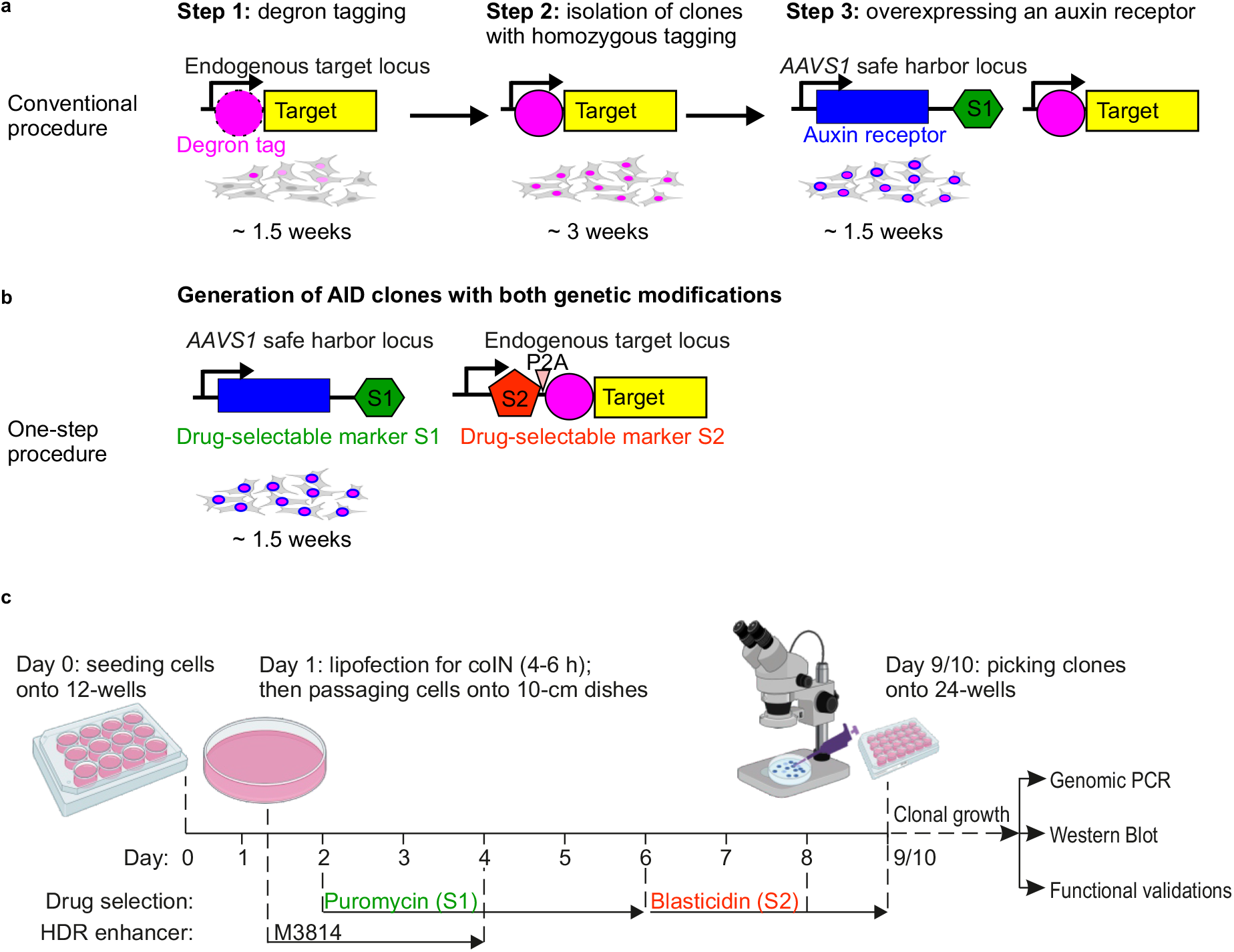
Overview of the conventional and one-step procedure to generate AID cells. **a-b,** illustration of genetic modifications and time required to generate AID clones with conventional procedure (a) and one-step procedure (b). P2A: self-cleaving peptide. **c**, timeline for one-step generation of AID clones in human cancer cell lines. Dashed line: timepoints for medium change or passaging/picking of cells.

Techniques to improve HDR efficiency and/or enrich HDR cells have been developed in recent years. These include use of HDR enhancers^17–20^, special design of HDR template^21,22^, 2A-drug selection cassette (2A: self-cleaving peptide)^23–25^ and co-incident insertion (coIN, or coselection) that additionally allows introduction of two genetic modifications in one step.^26,27^ In this work, we integrated the use of optimized AID components with coIN and other HDR enhancements, to achieve a rapid, single-step method for highly efficient generation of AID cells with homozygous degron-tagging. The procedure is scalable for multiple targets and applicable for several human cancer cell lines as well as hES cells, demonstrating the power of AID for dissecting the functions of endogenous proteins in diverse and differentiated human cells.

## Results

### Overview of one-step AID cell generation

Conventionally, AID cells are generated with low-efficiency homozygous tagging and require two engineering steps **(Fig. 1a)**.^8,10^ To streamline a one-step procedure with high efficiency, we took advantage of coIN and HDR enhancers to increase homozygous degron-tagging efficiency. Coordinated selection with puromycin (S1) and blasticidin (S2) enriched AID cells with both genetic modifications, i.e. degron tagging and auxin receptor expression **(Fig. 1b, c)**. Afterwards, manual picking under a stereo microscope enabled simple isolation of AID single-cell clones from 10-cm dishes **(Fig. 1c)**. The procedure is rapid, efficient and does not require other special equipment. It takes 1.5 weeks from initial cell seeding to clone isolation and only a small number of clones (typically 6-10) need to be screened. Simultaneous handling of 10-20 plates is feasible by a single operator. Below we describe the development of this procedure.

### Selection of AID components, coIN system and HDR enhancers

We first compared different options of the AID components, i.e. auxin receptor, degron tag and chemical inducer.^8,9,13^ We chose an AID system composed of *At*AFB2(F74A) as the auxin receptor, miniIAA7-3xFlag as the degron tag and 0.5 μM pico_cvxIAA as the inducer of proteolysis. The system uses a small degron tag, shows no basal degradation and achieves rapid inducible depletion with negligible off-target effects of the inducer **(Suppl. Fig. 1,** and **Suppl. Note 1**).

For efficient coIN, plasmids at a ratio of 1:3 (*AAVS1*: endogenous loci) were chosen to simultaneously introduce *At*AFB2(F74A) through HDR-mediated *AAVS1* safe harbor integration and tag endogenous loci with a degron **(Suppl. Fig. 2,** and **Suppl. Note 2)**. Testing of several HDR enhancers in coIN identified M3814^17,21^ and i53^18^ as effective HDR enhancers that increased degron-GFP tagging efficiencies **(Suppl. Fig. 3a-b,** and **Suppl. Note 2)**. Nearly 100% of cells were *At*AFB2(F74A)-mCherry positive after puromycin selection in all experiments if not specified (**Suppl. Fig. 2b**).

### HDR enhancers and coIN synergize to improve degron-tagging efficiency

We next tested 16 endogenous degron-GFP tagging pairs (8 templates with 2 different sgRNAs each) to assess the degron-tagging efficiencies through combining coIN and HDR enhancer M3814 (1 μM) in A431 cells. While coIN increased the percentage of GFP-positive cells by 1.5-fold, M3814 raised single-cell GFP intensities by about 30% **(Fig. 2a-c).** Higher single-cell GFP intensities with M3814 treatment indicate higher efficiencies of homozygous tagging, a desired property for generating AID cells. Strikingly, coIN and M3814 showed synergistic effects with an increase in both GFP-positive cells (2.3-fold) and single-cell GFP intensities (by 40%) **(Fig. 2a-c)**. M3814 additionally increased *At*AFB2(F79A)-mCherry levels **(Suppl. Fig. 3c)**, but showed considerable cytotoxicity, with about a 60% reduction in puromycin selected cells **(Fig. 2c)**.

**Figure 2:**
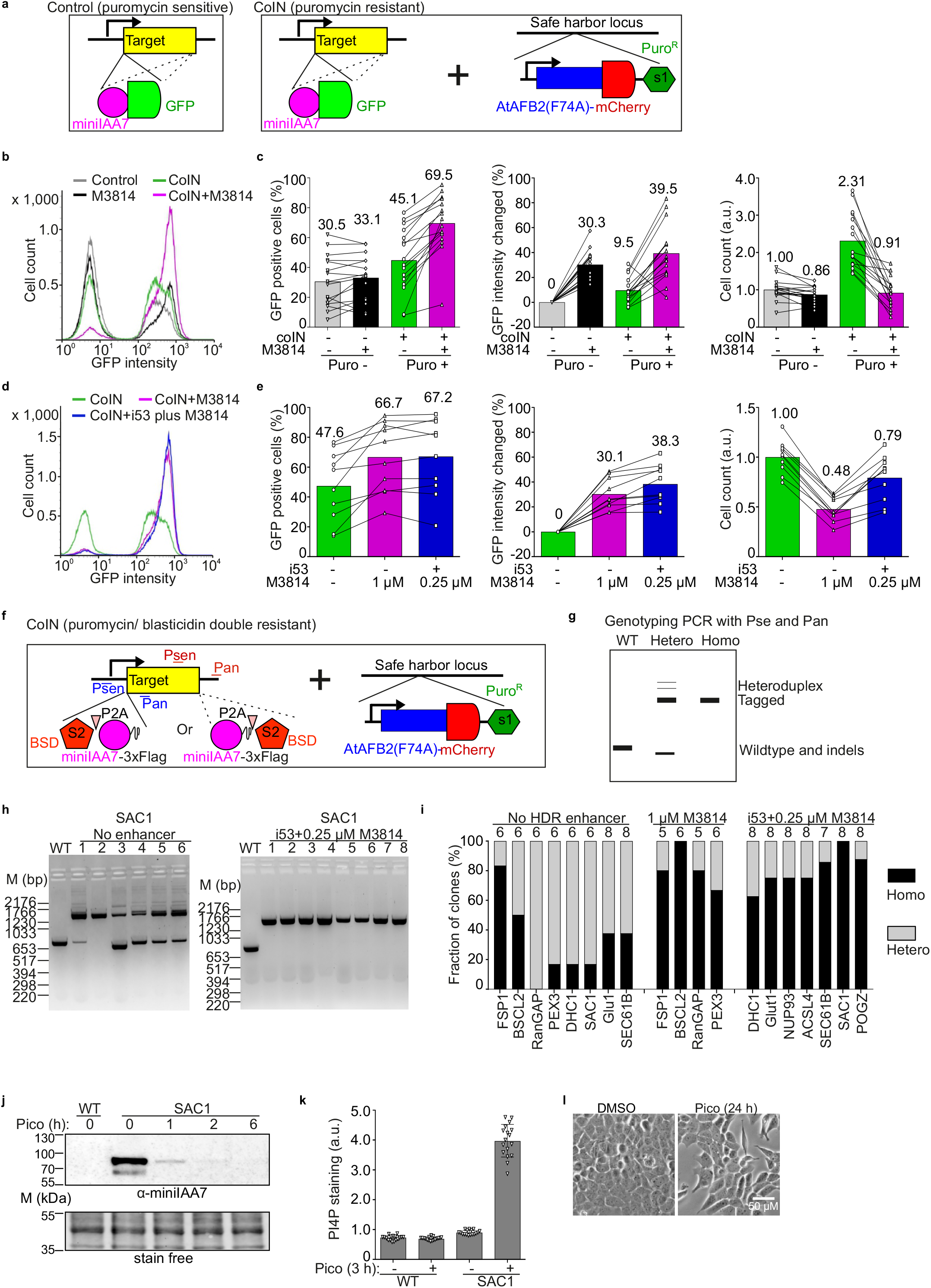
Optimization and applications of the one-step procedure in A431 cells. **a,** scheme of genetic modifications for degron-GFP tagging through conventional procedure (control) and one-step procedure (coIN) to assess tagging efficiencies in b-e. Puro^R^: puromycin-resistance gene. **b-c,** comparison of degron-GFP tagging efficiencies in conventional procedure and one-step procedure with 1 μM M3814 as HDR enhancer as indicated. A representative FACS profile (b) and statistics of percentage of GFP positive cells (c, left), single-cell GFP intensity change (c, middle) and cell count (c, right) are shown. N=16, 8 target proteins with 2 different sgRNAs each. **d-e,** comparison of degron-GFP tagging efficiencies in one-step procedure using different HDR enhancers. A representative FACS profile (d) and statistics of percentage of GFP positive cells (e, left), single-cell GFP intensity change (e, middle) and cell count (e, right) are shown. N=10, 5 target proteins with 2xsgRNAs each. Numbers above columns indicate mean values; lines link the same endogenous tagging pairs in (c, e). **f,** scheme of the genomic modifications in one-step generation of AID cells. BSD: Blasticidin S deaminase. Psen and Pan indicate the primer set for genotyping PCR of either N- or C-terminal tagging. **g,** scheme of a representative genotyping PCR result showing heterozygous and homozygous tagging at endogenous loci in the AID clones. **h,** genotyping PCR results of SAC1 clones generated with or without i53 plus 0.25 μM M3814 as HDR enhancers. **i,** statistics of genotyping PCR results for 8 targets without HDR enhancer, and 11 targets with either 1 μM M3814 or i53 plus 0.25 μM M3814 as HDR enhancers. Numbers above columns indicate total amount of clones analyzed. **j,** Western blot analysis of inducible SAC1 degradation. **k,** PI4P staining analysis upon SAC1 degradation. N=17 fields. **l,** wild-field imaging of cell morphological changes upon SAC1 degradation. Scale bar: 50 μM. Representative of 2 clones (j-k). WT: wildtype; Hetero: heterozygous; Homo: homozygous; pico: 0.5 μM pico_cvxIAA treatment; a.u. arbitrary unit.

I53 overexpression had low cell toxicity and an optional combination of i53 plus 0.25 μM M3814 as HDR enhancers was further evaluated (**Suppl. Fig. 3a-b)**. Testing of 10 degron-GFP tagging pairs demonstrated that i53 plus 0.25 μM M3814 outperformed 1 μM M3814 in cell counts and led to comparable increases in degron-GFP tagging efficiencies and *At*AFB2(F79A)-mCherry levels **(Fig. 2d-e,** and **Suppl. Fig. 3c)**. Collectively, these results demonstrate the outstanding performance of combined coIN and HDR enhancers to reach high homozygous tagging efficiencies in A431 cells in a single step.

### Establishment and characterization of AID clones in A431 cells

Two small size selection markers, Blasticidin S deaminase (BSD)^29^ and *Streptoalloteichus hindustanus* bleomycin (Sh_ble) genes^30^, were tested as 2A-drug selection cassettes to further improve degron tagging efficiencies. Results with Sh_ble are shown in **Suppl. Fig. 4a-b** and **Suppl. Note 3**. BSD had higher sensitivity than Sh_ble, effectively enriching all targets tested, and was thus used in the templates for endogenous tagging **(Fig. 2f**). AID clones were then isolated as outlined in **Fig. 1c**, with genotyping PCR (gPCR) to evaluate the homozygous tagging efficiencies in isolated clones **(Fig. 2g)**. Testing of 11 targets of widely different functions and expression levels ^8^ showed a degron-tagging efficiency of 100% (heterozygous plus homozygous) **(Fig. 2h-i)**. Clones generated with HDR enhancers (either 1 μM M3814 or i53 plus 0.25 μM M3814) for the 11 targets showed significantly higher homozygous degron-tagging efficiencies (average of 81%, varying from 62% to 100%) compared to the 8 targets without HDR enhancer (average of 32%, varying from 0% to 83%) **(Fig. 2h-i)**. Thus, HDR enhancers increased homozygous tagging efficiency in coIN, in line with the FACS analysis. Of note, RABGGTA tagged with miniIAA7-GFP without BSD selection achieved high-efficiency homozygous tagging of 90% with HDR enhancers i53 plus 0.25 μM M3814 (**Suppl. Fig. 4c-d**). Together, these results show that coIN with HDR enhancers achieves one-step generation of AID cells with high-efficiency homozygous tagging and that P2A-BSD effectively eliminates untagged cells.

The inducible degradation and ensuing functional readouts were then characterized for the 11 BSD targets and RABGGTA. A mouse monoclonal antibody against miniIAA7 (α-miniIAA7) was generated to facilitate the detection of degron-tagged proteins (see Methods). Based on Western blotting (WB), 11 of the target proteins were rapidly degraded in 1 h (**Suppl. Fig. 5a;** except for NUP93 that was effectively degraded after 6 h). NUP93 localizes in the nuclear pore complex and might have limited accessibility to *At*AFB2(F74A).^28^

Functional analysis of the targets revealed that upon pico_cvxIAA induction, most homozygous clones had expected phenotypic changes that were not observed in heterozygous clones (**Suppl. Fig. 5b-li)**.^29–31^ A good example is SAC1, the single known PI4P phosphatase in human cells that is required for cell viability.^32,33^ Rapid inducible degradation is thus optimal to study its functions. WB analysis showed that SAC1 was largely depleted after 1 h induction with pico_cvxIAA **(Fig. 2j),** accompanied by a 4-fold increase in anti-PI4P antibody staining intensity at 3 h induction **(Fig. 2k)** and a clearly altered cell morphology after 24 h induction **(Fig. 2l)**.

A few homozygous clones identified by gPCR exhibited reduced protein levels before induction and showed no clear functional readouts (**Suppl. Fig. 5a** and **d,** ACSL4_clone 1), potentially due to large deletions or other rearrangements in the target loci undetectable with gPCR.^17,34^ Use of 2-4 homozygous clones identified by gPCR is therefore recommended for the initial WB and functional analyses. Together, these results demonstrate rapid and effective degradation of the degron-tagged proteins in all the selected clones and excellent performance of *At*AFB2(F79A)/ miniIAA7-3xFlag/ pico_cvxIAA system for functional depletion of target proteins.

### Introduction of one-step AID system to other commonly used cell lines and analysis of clones

The established protocol was next tested in other widely used human cell lines, including lung alveolar cancer A549, embryonic kidney HEK293A, osteosarcoma U2OS and prostate cancer derived PC3 cells. CoIN of 6 endogenous degron-GFP tagging pairs (3 templates with 2 different sgRNAs each) was first performed with or without HDR enhancers. PC3 cells died out after puromycin selection, likely due to deficiency of the HDR repair pathway.^35^ In the other 3 cell lines, effective degron-GFP tagging was achieved using coIN and further improved with HDR enhancers as in A431 cells **(Fig. 3a-f).** Substantially lower degron-GFP tagging efficiency was obtained through conventional tagging without coIN and HDR enhancer **(Suppl. Fig. 6a)**. In these 3 lines, i53 plus 0.25 μM M3814 again outperformed 1 μM M3814, yielding similar enhancement of HDR efficiency with lower cytotoxicity **(Fig. 3a-f)**. With HDR enhancers, the expression of *At*AFB2(F79A)-mCherry was again slightly increased in all 3 cell lines **(Suppl. Fig. 6b).** These results indicate that the protocol established works similarly for high-efficiency homozygous degron-tagging in several human cancer cell lines proficient in HDR.

**Figure 3:**
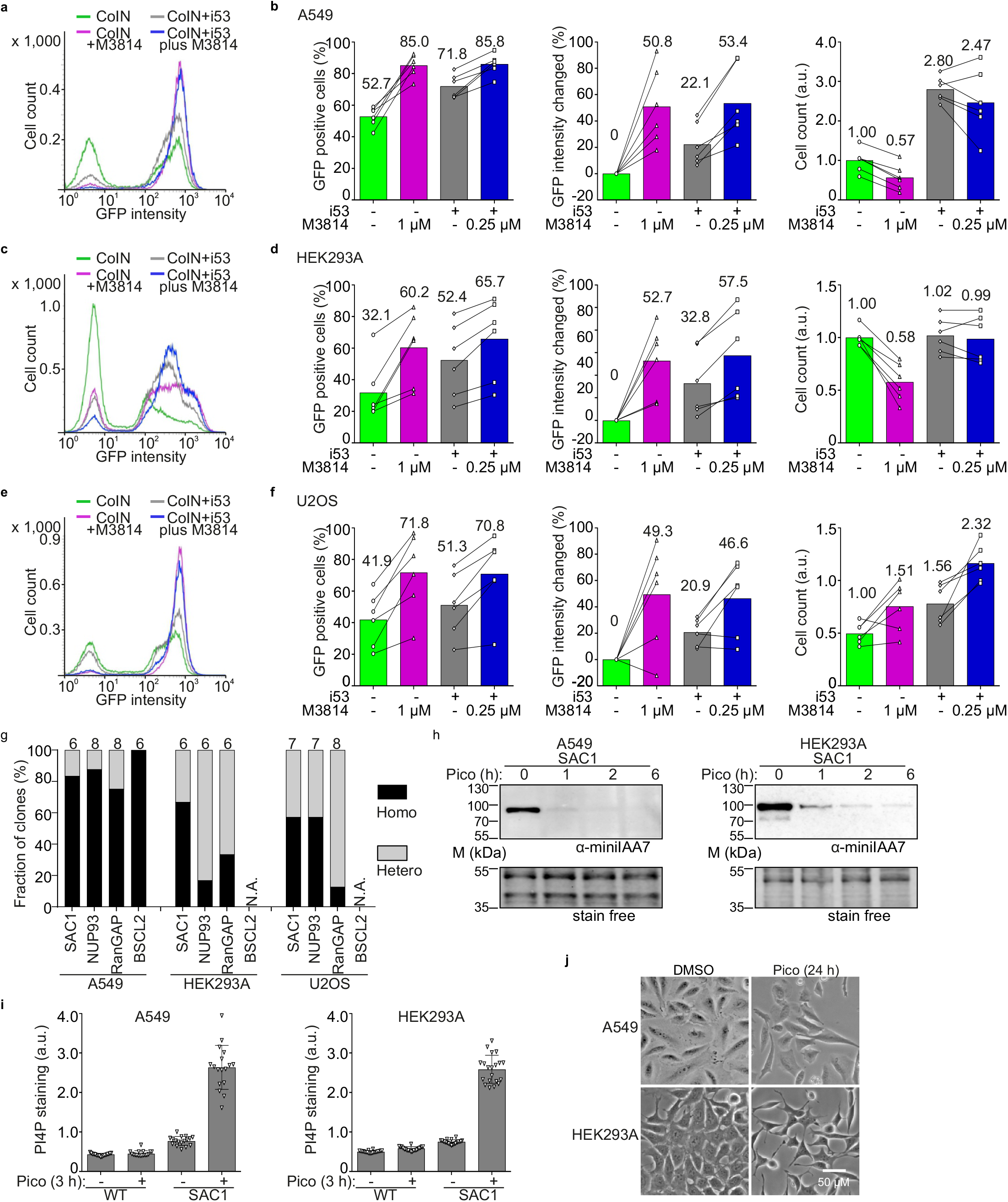
Evaluation and applications of the one-step procedure in other human cancer cell lines. **a-f,** comparison of degron-GFP tagging efficiencies in coIN with different HDR enhancers in A549 (a-b), HEK293A (c-d) and U2OS (e-f) cells. Representative FACS profile (a, c, e) and statistics of percentage of GFP positive cells, single-cell GFP intensity change, and cell count (b, d, f) are shown. Numbers above columns indicate mean values; lines link the same endogenous tagging pairs. N=6, 3 protein targets with 2xsgRNAs each. **g,** statistics of genotyping PCR results for 4 targets with i53 plus 0.25 μM M3814 as HDR enhancers. Clones were generated and identified as indicated in Fig. 2f-g. Total number of clones analyzed is indicated above each column. N.A.: not available due to cell death after selection with blasticidin (S2). **h,** Western blot analysis of inducible SAC1 degradation. **i,** PI4P staining analysis upon SAC1 degradation. N=17 (A549) and 20 (HEK293A) fields. **j,** wild-field imaging of cell morphological changes upon SAC1 degradation. Scale bar: 50μM. Representative of 1 (A549) and 2 (HEK293) clones (h-j). WT: wildtype; pico: 0.5 μM pico_cvxIAA treatment; a.u.: arbitrary unit.

Single-cell clones were then generated as for A431 cells **(Fig.1 c** and **Fig. 2f-g)**. For A549 and U2OS cells, clones were isolated directly from 10-cm plates. For HEK293A cells, limited dilution cloning into 96-well plates was used as the cells migrated extensively during growth and formed clones without a clear boundary. Of the 3 cell lines, A549 showed the best homozygous degron-tagging efficiency **(Fig. 3g).** BSCL2 clones failed to be isolated in HEK293A and U2OS cells, as the clones died after blasticidin (S2) selection, indicating that the cells might express Seipin/BSCL2 at lower levels or be more sensitive to blasticidin **(Fig. 3g)**. Further improvement in the selection (for example by using a more sensitive selection marker as S2) might solve the problem. Alternatively, single clones can be isolated without S2 selection as for RABGGTA in A431 cells **(Suppl. Fig. 4c-d)**.

Regarding functional effects, WB and microscopy analyses of SAC1 degron A549 and HEK293A cells showed rapid protein degradation, expected increase of cellular PI4P and morphological changes analogous to those observed in A431 cells **(Fig. 3h-j).** Similar changes as in A431 cells were also found for the other target proteins tested in A549 and HEK293A cells, including WB and phenotypic readouts **(Suppl. Fig. 7a-d** and **f).** In general, U2OS cells exhibited a somewhat slower degradation of all 3 target proteins post-induction, despite proper expression of *At*AFB2(F79A)-mCherry **(Suppl. Fig. 7a** and **e).** The compromised inducible protein degradation in U2OS cells might indicate a lower activity of other SCF E3 ligase components in these cells.

### One-step introduction of AID system to H9 hES cells

Homozygous tagging is of low efficiency in hES or iPS cells.^15,16^ Accordingly, our early attempts to tag endogenous POGZ locus in H9 cells using electroporation resulted in only 0.1% homozygously tagged cells **(Suppl. Fig. 8a-c)**. Moreover, the expression of *At*AFB2(F74A) under the EF1a promoter was unstable in H9 cells, leading to inefficient inducible degradation **(Suppl. Fig. 8d-e)**. Further optimization revealed that the use of CAG promoter improves both stable expression of *At*AFB2(F79A)-mCherry (from 50-60% mCherry positive cells with EF1a promoter to 99% with CAG promoter) and *AAVS1* integration efficiency (by about 2.5-fold compared to CMV promoter to express Cas9) **(Fig. 4a-b)**. These results emphasize the importance of promoter choice for successful establishment of AID cell lines in human stem cells.^36^

**Figure 4:**
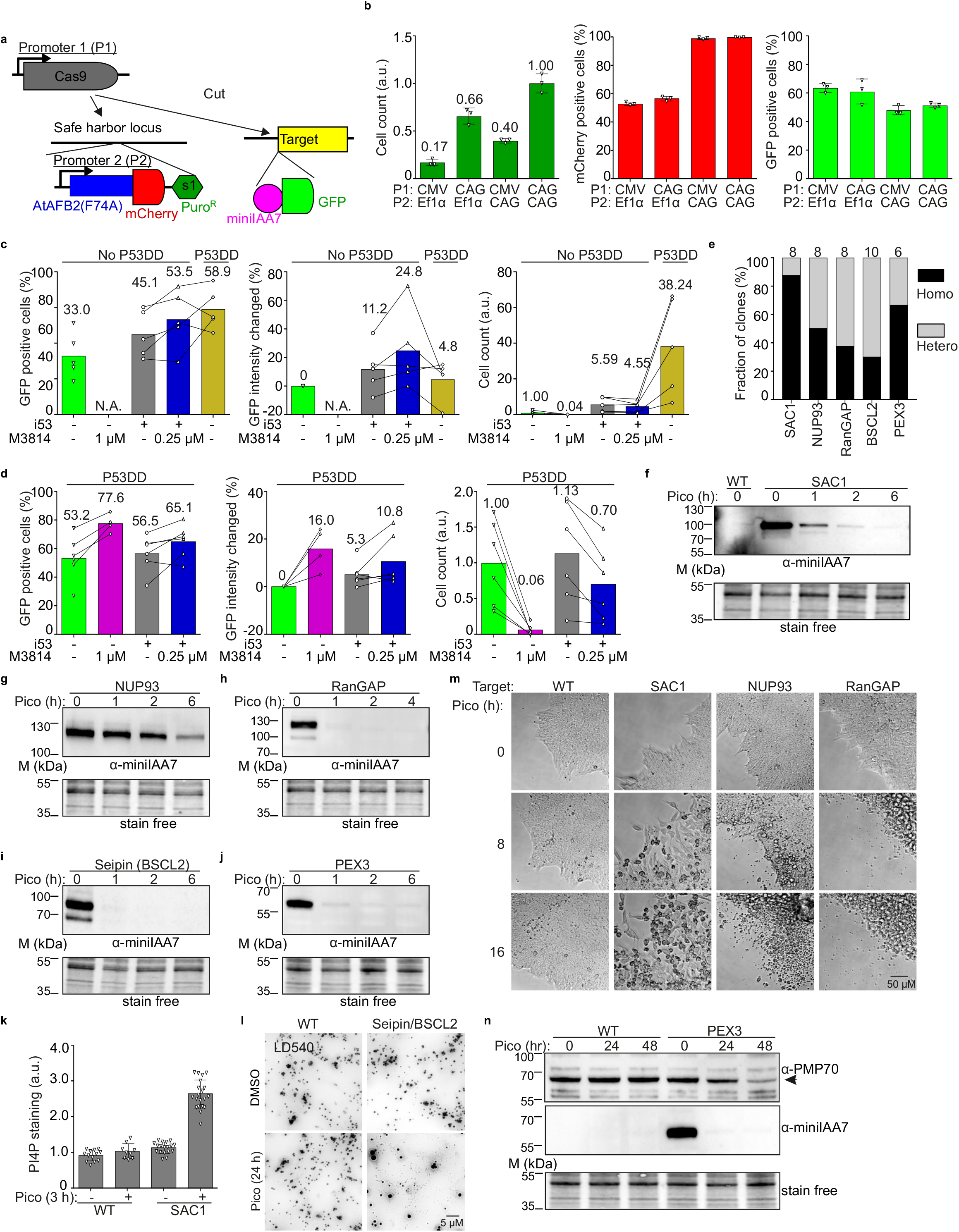
Optimization and applications of the one-step procedure in H9 human embryonic stem cells. **a,** scheme illustrating the two promoters P1 and P2 in plasmids for one-step generation of AID cells. **b,** comparison of cell count, fraction of mCherry positive and GFP positive cells using different P1 and P2 promoters in H9 hES cells. Stable cell pools after puromycin selection were used for the analysis. N=3 technical repeats. **c-d,** statistics of percentage of GFP positive cells, single-cell GFP intensity change and cell count with different HDR enhancers with or without P53DD. P53DD: a dominant negative P53 mutant; numbers above columns indicate mean values; lines link the same endogenous tagging pairs. N.A.: not available due to insufficient number of cells for FACS analysis. N=5 (c) and 6 (d) different protein targets. **e,** statistics of genotyping PCR results for 5 targets in H9 cells with i53 plus 0.25 μM M3814 as HDR enhancers in the absence of P53DD. Clones were generated as indicated in Fig. 2f-g. Total amount of clones analyzed is indicated above each column. **f-j,** Western blot analysis of inducible degradation of 5 target proteins in H9 degron cell lines with anti-miniIAA7 antibody. **k,** PI4P staining analysis of H9 wildtype and SAC1 degron cells. N=13, 9, 18, 20 fields for the 4 conditions. **l,** lipid droplet (LD) staining with LD540 in wildtype and BSCL2/Seipin degron H9 cells. 0.2 mM of oleic aicd (OA) was added during the final 4 h to induce LD formation. Scale bar: 5 μM. **m,** live-cell imaging analysis of morphological changes and cell death in H9 wildtype and 3 different degron cells. Representative images in different time points from the same areas are shown. Scale bar: 50 μM; N=4 fields. **n,** Western blot analysis with anti-PMP70 and anti-miniIAA7 antibodies in H9 wildtype and PEX3 degron cells. Arrow indicates the specific PMP70 protein bands. Representative of 2 clones for each target protein (f-n). WT: wildtype; pico: 0.5 μM pico_cvxIAA treatment; a.u. arbitrary unit.

Using these optimizations, we tested degron-GFP tagging pairs for 5 different targets using coIN and HDR enhancers as well as P53DD, a dominant negative P53 mutant. P53DD was shown to substantially improve the viability of engineered human stem cells that are sensitive to Cas9 induced double-strand breaks (DSBs) in a P53-dependent pathway.^37^ In coIN, P53DD dramatically increased puromycin resistant cell counts by roughly 40-fold, and the percentage of degron-tagging cells almost 2-fold without a clear impact on single-cell GFP intensity **(Fig. 4c)**. Interestingly, i53 increased the cell count by 5-fold in the absence of P53DD, but not in its presence **(Fig. 4c-d)**. The results imply that i53 might partly inhibit a P53-dependent pathway to improve cell viability. M3814 at 1μM showed severe cytotoxicity that reduced cell counts by about 20-fold and was not rescued by P53DD (**Fig. 4c-d).** Instead, i53 plus 0.25 μM M3814 increased the efficiency of degron-tagging, single-cell GFP intensity and cell count, with P53DD inhibition further increasing cell counts **(Fig. 4c-d).** Finally, one-step generation of AID clones with i53 plus 0.25 μM M3814 (here without P53DD) allowed the isolation of H9 single-cell clones in 1.5 weeks and showed an average homozygous degron-tagging efficiency of >50% (ranging from 30-87%) for 5 targets **(Fig. 4e).**

### Protein degradation in H9 hES cells

WB analysis showed that 4 of the targets were degraded in 1 h and NUP93 was again degraded more slowly (in about 6 h) of pico_cvxIAA treatment **(Fig. 4f-j)**. Functional analysis revealed expected phenotypes for all the targets: rapid increase of PI4P staining in SAC1 degron cells (3 h induction), lipid droplet biogenesis defects in BSCL2/seipin degron cells (24 h induction), and extensive degradation of peroxisomal membrane protein PMP70 in PEX3 degron cells (24-48 h induction) **(Fig. 4k, l** and **n).** Furthermore, live-cell imaging showed specific and distinct morphological changes preceding cell death in SAC1, NUP93 and RANGAP1 degron cells within 16 h of induction **(Fig. 4m).** WB with Oct3/4 antibody indicated that the isolated degron cell lines maintained their polypotency **(Suppl. Fig. 8f).** Together, these results demonstrate that AID is powerful for chemogenetic control of endogenous proteins in H9 hES cells.

### Chemogenetic control of endogenous protein degradation in H9-derived neurons

H9 cells have the potential to differentiate into other cell types. To study if H9 AID cells can first be differentiated and protein degradation then be induced only at the differentiated cell stage, we differentiated H9 AID cells to neurons with an established and highly efficient procedure.^37^ Morphological and WB analysis with the neuronal marker anti-β-Tubulin III confirmed the successful neuronal differentiation of the AID cell lines **(Fig 5a-d, g-h).** Upon pico_cvxIAA induction, rapid and efficient depletion of the target proteins was achieved in differentiated cells, as shown by WB (in 1 h for SAC1, PEX3 and RANGAP1; in 6 h for NUP93). This was followed by expected functional readouts, i.e. an increase of PI4P staining in SAC1 degron cells, degradation of PMP70 in PEX3 degron cells **(Fig. 5e-g)** and cell death in RANGAP1 and NUP93 degron cells **(Fig. 5h).** These findings show the applications of AID for chemogenetic control of endogenous proteins in hES-derived neurons and demonstrate the power of AID in human stem cell biology to increase our understanding of protein functions in differentiated cell types.

**Figure 5:**
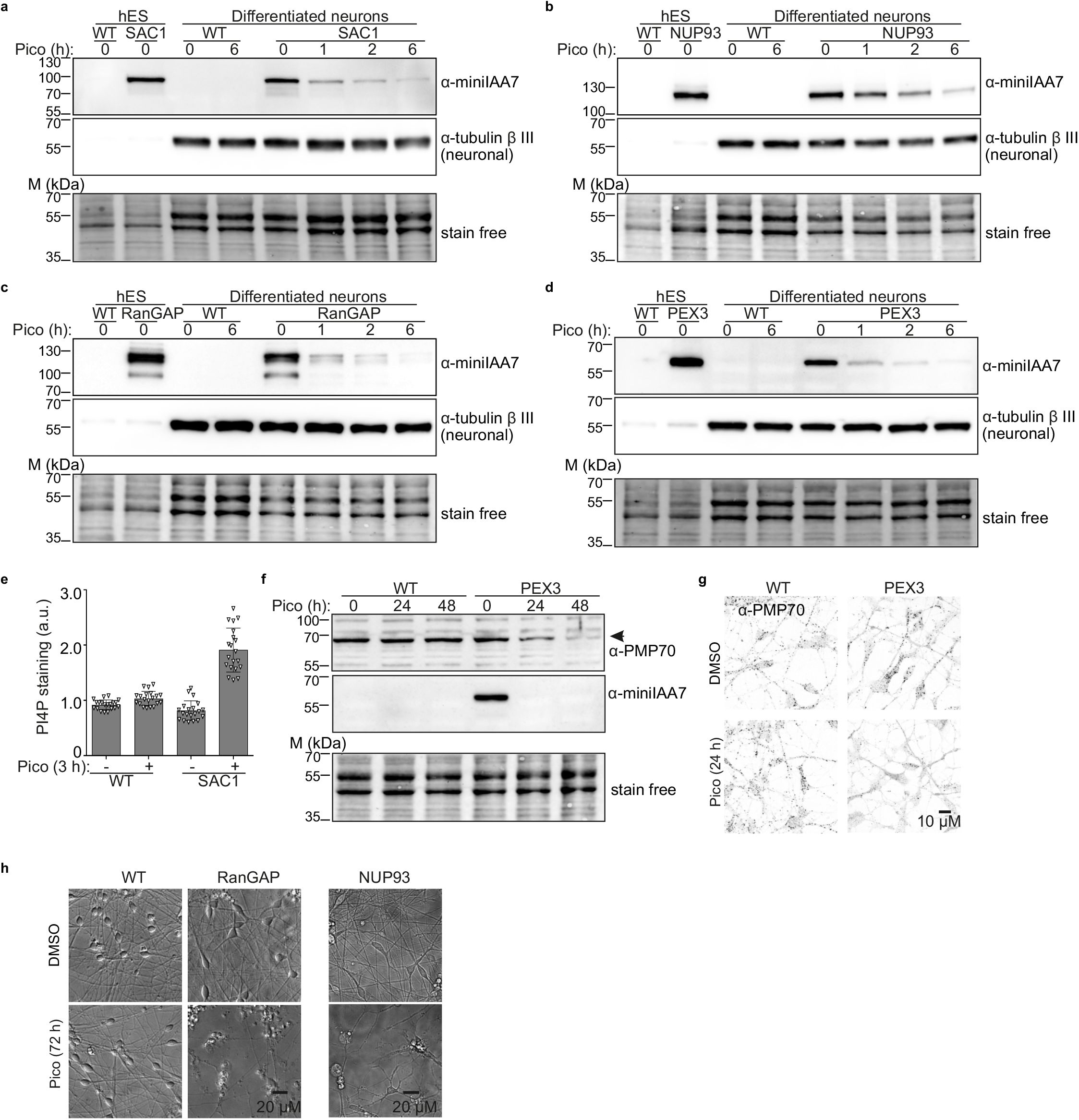
Chemogenetic control of endogenous protein degradation in neurons differentiated from H9 cells. **a-d,** Western blot analysis in neurons differentiated from H9 degron cells targeting SAC1 (a), NUP93 (b), RANGAP1 (c) and PEX3 (d). **e,** PI4P staining analysis in neurons differentiated from wildtype and SAC1 degron H9 cells. a.u. arbitrary unit. N=20 fields. **f-g,** Western blot analysis (f) and immunofluorescence staining (g) of peroxisomal membrane protein PMP70 in neurons differentiated from wildtype and PEX3 degron H9 cells. Scale bar: 10 μM. **h,** live-cell imaging of cell death in neurons differentiated from wildtype, RANGAP1 and NUP93 degron H9 cells. Scale bar: 20 μM. Representative of 1 (RANGAP1 and NUP93) and 2 (SAC1 and PEX3) cell clones. WT: wildtype; pico: 0.5 μM pico_cvxIAA treatment.

## Discussion

Loss-of-function analysis is one of the most important strategies for understanding protein functions in mammalian cells.^38^ AID provides a powerful means to achieve this in a rapid, inducible manner but requires the introduction of two genetic modifications. So far, efforts for one-step generation of AID cells have employed a rescue strategy through CRISPR/Cas9-mediated knockout in conjunction with the expression of a degron-tagged rescue construct plus an auxin receptor.^39–41^ This strategy may suffer from downsides of CRISPR knockout, including compensations that can rescue target activities.^42,43^ Critically, single clones show considerable variations of target protein expression as the rescue constructs are randomly inserted into the genome at various copy numbers.^39–41^ The rescued cells further suffer from lack of endogenous transcriptional and translational control, such as response to environmental stimuli or cell differentiation cues.^38–40^ Intensive identification of single clones was thus needed, and applications were biased towards essential target proteins.^39–41^

Here, we harnessed recent developments of the AID technology to optimize its performance **(Suppl. Fig. 1).**^8,9,11,13^ We then established a procedure for one-step generation of AID cell clones through endogenous degron-tagging with an unprecedented speed of 1.5 weeks **(Fig. 1c)** and tagging efficiencies of 100% in the isolated clones (homozygous plus heterozygous) for multiple target proteins in 5 cell lines **(Fig 2i, 3g** and **4e).** Extensive isolation and identification of single clones was not required since 1) the tagged proteins were expressed under the control of endogenous promoters and 2) high-efficiency homozygous tagging was achieved with near 100% expression of the auxin receptor.

Somewhat surprisingly, we found that HDR enhancers substantially improved homozygous tagging efficiency. Moreover, HDR enhancer and coIN synergistically improved the degron-tagging efficiency while coIN achieved one-step generation of AID cells. Yet, the cytotoxicity of HDR enhancers needs to be considered. The most proficient HDR enhancer M3814 clearly reduced cell counts, especially in the sensitive H9 cells. Use of i53 reduced the concentration of M3814 required to achieve high tagging efficiency, resulting in higher cell counts in all the cell lines studied. Interestingly, i53 additionally increased the number of A549 and H9 cells compared to control, possibly via inhibiting P53-dependent cell death caused by Cas9-induced DSBs **(Fig. 3b and 4c-d)**.^44^

The importance of homozygous tagging is best illustrated by functional consequences upon induction of degradation. Heterozygously tagged cells often manifested no clear phenotypic changes **(Supplementary Fig. 5d, e** and **g)**. Of note, gPCR cannot detect clones with a long deletion, leading to false identification of heterozygously tagged clones with a long deletion as homozygously tagged ones.^17,34^ These clones showed reduced degron-tagged target levels and compromised functional readouts **(Supplementary Fig. 5,** ASCL4_clone 1; and **Supplementary Fig. 7,** Seipin_clone 2). Thus, it is advisable to select several homozygous clones identified with gPCR for the initial Western blot analysis and functional tests.

The choice of cell line has a major impact on the established procedure. Cells deficient in HDR, such as PC3, would not be suitable for it.^35^ Of the cell lines tested, A431 and A549 cells were the most proficient. While other cell lines were successfully engineered, HEK293A, U2OS and H9 cells displayed somewhat lower homozygous tagging efficiencies. Moreover, cells that do not grow in colonies (such as HEK293) could be isolated by limiting dilution, and more slowly proliferating cells (such as A549, U2OS) need a longer time for AID clone isolation. To apply the procedure in a new cell line, the *AAVS1* integration plasmid pairs and/or SEC61B endogenous tagging plasmid pairs could be used to rapidly assess whether the cell line is HDR proficient. For cell lines that are deficient in HDR, recent development in new genetic engineering tools, such as PASTE^45^ that combines prime editing^46^ with site-specific integrase^47^, might enable homozygous tagging of endogenous proteins.

For H9 cells, additional optimizations were required regarding 1) the promoter to enable uniform auxin receptor expression and 2) use of i53 and/or P53DD to mitigate cytotoxicity **(Fig. 6a-d)**. With these improvements, we successfully isolated single cell clones for the 5 targets reported. Of note, H9 clones with mosaic expression were observed in some other attempts, possibly due to inefficient single cell dissociation with accutase and/or the rapid proliferation of H9 cells. A further step of single-cell cloning after generation of AID cell pools could be applied in such cases.

The introduction of AID for chemogenetic control of endogenous proteins in human ES cells holds great promise in stem cell biology. Chemical induction is favorable for stem cells as it is non-invasive, rapid, efficient, and flexible.^48^ Moreover, differentiated cell types are refractory to genetic modification and not proficient in HDR.^49^ We demonstrated AID of several endogenous targets in hES cells with high efficiency and provided the first evidence that AID removes endogenous proteins in cells differentiated from them, using neurons as a test case. In the future, genetic engineering in human stem cells in combination with cell differentiation procedures would allow chemogenetic degradation of endogenous proteins in various cell types and opens entirely new possibilities for e.g. disease modeling and cell therapy.

## Supporting information

LiS et al. 2023 AID supplementary data

## Acknowledgements

We thank A. Uro and R. Kosonen for technical assistance. HiLIFE Flow cytometry and Light microscopy core facilities are acknowledged for access to research infrastructure, Wales Gene Park for performing NGS and the Supercomputing Wales project, which is part-funded by the European Regional Development Fund (ERDF) via Welsh Government. Fig. 1a and c were created with templates from BioRender. This study was supported by Academy of Finland (grants 324929 to EI, 322647 to VMO, 332096 to XZ), Jane and Aatos Erkko Foundation (EI), Foundation Leducq (grant 19CVD04 to EI), Sigrid Juselius Foundation (EI, VMO), EU MSCA-PF (grant 101059424 to MvdS), The Hospital District of Helsinki and Uusimaa/Children and Adolescents (TR), The Paulo Foundation (N.E.), The Maud Kuistila Memorial Foundation (N.E.), and H2020-MSCA-IF (grants 894596 to YW, 813707 to SM).

## Author contributions

SL and EI conceptualized the project with assistance from YW (hES). SL designed and performed most of the experiments with assistance from YW (hES culture and engineering), MvdS (gPCR, WB, hES engineering, LD staining), XZ (PI4P experiments), KK (PMP70 staining), VDN and YZ (RNAseq). YW designed and performed neuron differentiation with assistance from SM. NE helped with POGZ tagging in hES. VMO provided resources. TR supervised the hES work by YW, SM and NE. EI acquired funding and supervised the study. SL and EI wrote the manuscript with help from MvdS and other coauthors.

## Notes

### Competing Interest Statement

The authors have declared no competing interest.

