## Supplementary material for "One-step generation of auxin-inducible degron cells with high-efficiency homozygous tagging": LiS et al. 2023 AID supplementary data

### Methods:

#### Culture and transfection of human cancer cell lines

A431 cells (ATCC CRL-1555), U2OS cells (kindly provided by Marikki Laiho at John Hopkins University, USA) and HEK293A cells (Invitrogen R70507) were cultured in DMEM (high-glucose, Lonza/Gibco), and A549 cells (ATCC CCL-185) in F-12 Nutrient Mixture (Gibco), supplemented with 10% FBS, penicillin/streptomycin (100 U $ml^{-1}$ each), L-glutamine (2 mM) at 37 °C in 5% $CO_2$. Cells were tested negative for Mycoplasma using PCR detection. Cells were seeded for 16 h in 12-well and transfected at 80–95% confluence using Lipofectamine LTX with PLUS Reagent (Invitrogen 15338-100), typically with 1.0 µg plasmid(s) per 1.0 µl of PLUS reagent (1.5 µl for HEK293A), 2.0 µl of Lipofectamine LTX (3.0 µl for HEK293A) and $4.0 \times 10^5$ (A431 and HEK293A) or $3.0 \times 10^5$ (U2OS and A549) cells per well.

#### Chemicals and antibodies

Nedisertib (M3814, Selleckchem S8586) prepared as 10 mM stock in DMSO; cvxIAA (TCI M3141) 25 mM stock in DMSO, 5-Adamantyl-IAA (pico_cvxIAA, TCI A3390) 2.5 mM diluted from 100 mM stock in DMSO, 5-PH IAA (MCE HY-134653) 20 mM in DMSO, Indole-3-acetic acid sodium (IAA, Sigma I5148) 0.25 M stock in $H_2O$, puromycin (Sigma, P8833) 10 mg/ml in $H_2O$, Blasticidin S HCl (Gibco A1113903) 10 mg/mL solution, Zeocin (Gibco R25001) 100 mg/ml solution, LD540 (synthesized by Princeton BioMolecular Research) 1 mg/ml in Ethanol, Erastin (Selleckchem S7242) 10 mM in DMSO, Oleic Acid (OA, Sigma O-1383) 1 mM as OA:BSA complex (8:1 molar ratio) in serum free DMEM, goat anti-FABP4 (WB 1:500, Santa-Cruz sc-18661), mouse monoclonal anti-tubulin beta III (neuronal) (WB 1:2,000 Sigma, T8578-100UL), mouse monoclonal anti-PMP70 (WB 1:1,000, Sigma-Aldrich SAB4200181), mouse monoclonal anti-PI4P IgM (IF 1:200, Echelon Z-P004), mouse monoclonal anti-Oct3/4 (WB 1:500, Santa Cruz sc-5279), goat anti-mouse IgG (H/L) HRP (1:1,000, Bio-Rad 1706516), donkey anti-goat (H/L) IgG HRP (1:10,000, Jackson immuno research 705-035-147), goat anti-mouse IgM (H) Alexa Fluor 488 (1:200, Thermo Fisher A21042).Mouse anti-miniIAA7 (WB final 0.5 µg/ml, Ximbio 158027) was generated by Genscript. Briefly, the 20 aa. core peptide of miniIAA7 tag with an extra C conjugated to KHL (QVVGWPPVRNYRKNMMTQQKC-KHL) was used as the immunogen and BALB/c mouse was used as the host strain. Antisera from 5 mice were tested in WB with endogenous miniIAA7 tagged A431 cell lysates. The best one was chosen for cell fusion to generate hybridoma cell clones. The clones were screened again with WB and 5 of the best clones were selected for subcloning to get the final clone 12A8-1. 12A8-1 was used for antibody production to generate purified mouse IgG and designated as mouse anti-miniIAA7 (12A8-1).

#### Construction of plasmids

Vector sequences are provided in **Supplementary sequences**.

To express different auxin receptors through *AAVS1* integration, pSH-EFIRES-P-*At*AFB2-mCherry (Addgene 129716) was used as a backbone. *Os*TIR1 was synthesized by Genscript as codon-optimized cDNA and substituted *At*AFB2 on the backbone through restriction ligation. Overlap PCR was used to introduce F74A and F74G point mutations in the auxin receptors. CAG promoter was cloned from plasmid AAS1815 (Addgene 107942)[50] to substitute EF1a promoter on the backbone where indicated.

To express Cas9 and different sgRNAs, sgRNAs were synthesized as two unphosphorylated primers, annealed and inserted into BbsI-cut pCas9-sgRNA (as Addgene 129726) or pCas9/VRQR-sgRNA (Addgene 129725) vectors. CAG promoter from plasmid AAS1815 (Addgene 107942) was used to substitute the CMV promoter on the vectors to express Cas9 where indicated. SgRNA targeting sites were searched manually for –NGG PAM sequence within 18 bp after insertion sites or CCN- within 18 bp before insertion sites. The sgRNA target sites were disrupted in the templates by the insertions.

To construct HDR templates of endogenous targets, homology arms on donor vectors were amplified from A431 genomic DNA through PCR using Q5 High-Fidelity DNA Polymerase with High GC Enhancer (NEB, M0491). Tags were synthesized as codon-optimized cDNA and cloned into the donor through restriction ligation or overlap PCR. The PCR fragments were cloned into pGL3-basic backbone using NEBuilder HiFi DNA Assembly Master Mix (NEB, E2621) or through restriction ligation. Tags on the HDR templates were changed through restriction ligation or Gibson assembly with NEBuilder HiFi DNA Assembly Master Mix.

I53 and P53DD were synthesized as codon-optimized cDNAs by Genscript and cloned into pGL3-basic vector with EF1a promoter and HSV TK poly(A).

**A431 cell lines for testing different auxin receptors**

pSH-IRES-B-Seipin-miniIAA7-mEGFP (Addgene 129719) was used to generate single clones expressing Seipin-miniIAA7-mEGFP through HDR-mediated *AAVS1* integration. A431 single clones with homozygously tagged DHC1-miniIAA7-mEGFP, Seipin-miniIAA7-mEGFP, and heterozygously tagged EGFR-miniIAA7-mEGFP have been described.[8] To express different auxin receptors, *AAVS1* integration vector expressing auxin receptors (0.6 µg) and pCas9-sgAAVS1-1 (0.4 µg, Addgene 129726) were co-transfected into cells with Lipofectamine LTX with PLUS Reagent for 24 h, followed by selection with 1 µg/ml puromycin for 6 days. Cell pools were used for FACS and live-cell imaging.

**One-step procedure for generation of human AID clones in cancer cell lines**

An overview of the procedure is provided at **Fig 1c.** A431, U2OS, HEK293A and A549 cells were seeded on 12-well plate at day 0 and transfected with a mixture of 4 plasmids (*AAVS1*: target at 1:3 ratio as shown in **Suppl. Fig 2e**) or 5 plasmids (0.8 µg of 4 plasmids plus 0.2 µg i53 plasmid) at day 1 using Lipofectamine LTX with PLUS Reagent. After 4-6 h, one third of transfected cells was passaged to a 10 cm dish containing the indicated concentration of M3814 (0, 0.25 or 1 µM). At day 2, medium was replaced with fresh medium containing 1 µg/ml of puromycin and the same concentration of M3814 as in day 1. At day 4, fresh medium with puromycin but without M3814 was used. At day 6 and 8, fresh medium with 10 µg/ml Blasticidin was used to select clones with endogenous tagging.

At day 9-10, single clones formed on the 10-cm plates. Picking of single clones is analogous to iPS clone picking with videos available online. The picking skill can generally be learned on the first attempt and is easy to master. Briefly, a S9 E StereoZoom microscope (Leica, 10450814) with 10x Eyepieces on a TL3000 ergo light base (Leica, 10450690) was set up to check and isolate single clones. Before picking of single clones, medium on the 10-cm plates were changed to PBS or antibiotic-free medium to help the survival of clones after picking. A 24-well plate with regular medium was prepared to grow the clones. Clones formed on the 10-cm plate can be visualized on the TL3000 ergo light source with proper contrast. Clones on the plate were moved to the centre, checked under the objectives, and picked with a regular 10 or 20 µl pipette. The pipette with a tip was gently pressed beforehand and the single clones were detached by the pipette tip with gentle mechanical scraping. When cells were detached, the pressed pipette was released slowly to suck in the detached cells and the cells were transferred to 24-well plate with regular growth medium. 2-3 days later, clones on the 24-well plates can be passaged with trypsin for further expansion and characterization. To be noted, single clones on the 10-cm plate are clearly faster growing than on a 96-well plate, as on the 10-cm plate gas exchange and cell-cell communication are maintained better,and medium can be simply changed to improve the growth of cells. Moreover, clone densities on the 10-cm plate are flexible and densities of 1-1,000 clones per plate can be easily isolated.

**FACS analysis**

Cells were seeded at 1:5 (for A431 and HEK293A) or 1:3 (for A549 and U2OS) into a six-well plate. On day 1, medium was changed to 2 ml fresh medium without (for 0 h and 1 h induction) or with (for 16 h induction) inducers. On day 2, the 1 h samples were supplemented with 0.5 ml medium containing 5x concentration of the indicated inducer and incubated for 1 h at 37 °C. After treatment, cells were detached with 0.5 ml trypsin at 37°C for 5–8 min (U2OS, HEK293A and A549) or 8–12 min (A431), put on ice and transferred to 1.5 ml Eppendorf tubes containing 0.5 ml serum-free $CO_2$ independent medium (Gibco 18045088). The cell suspensions were centrifuged at 4 °C, resuspended in 0.3 ml ice-cold serum-free $CO_2$ independent medium and stored on ice before FACS analysis. FACS analysis was performed on a BD Influx cell sorter (BD Biosciences) with a 100 µm nozzle at 4–8 °C using BD FACS Software. Cells were gated with SSC, FSC and trigger pulse width for singlets, and 50,000-100,000 cells were analysed from each sample. GFP was excited with a 488 nm laser and detected with a 530/40 detector; mCherry was excited with a 561 nm laser and detected with a 615/20 detector. Data were analysed with BD FACS Software. Background subtracted mean fluorescence intensity was used for analysis.

**Cell counting**

Cells were counted with Bio-Rad TC10 automated cell counter (Bio-Rad 145-0001) using 10-20 µl of cell suspension in TC10 counting slides (Bio-Rad 145-0015). Histograms of cell diameter distribution were checked after each count to avoid counts with abnormal histograms.

**Reversibility assays**

For inducer washout experiments, cells were seeded at day 1 on µ-slide 8-well ibiTreat dishes. On the second day, cells were treated with the indicated inducers overnight. On the third day, cells were washed 4 times with FluroBrite DMEM containing 10% FBS without inducer before live-cell imaging. Cells were imaged immediately after washing with Nikon Eclipse Ti-E widefield microscope equipped with 20x air objective NA 0.8, Nikon Perfect Focus System 3, Hamamatsu Flash 4.0 V2 scientific CMOS and Okolab stage top incubator system. Multipoint and time lapse imaging was started immediately and recording was every 30 min for 18 h. Background subtracted fluorescence intensities were used for analysis.

**RNAseq and analysis**

RNA samples were extracted using NucleoSpin RNA Mini kit (MACHEREY-NAGEL 740955). For RNA sequencing, library was prepared using NEB Ultra II directional RNA library prep kit (NEB E7760). Samples were sequenced using an Illumina HiSeq 4000 with 75 bp paired-end reads. Sequencing reads of all samples were first quality controlled using FastQC v0.11.8 (http://www.bioinformatics.babraham.ac.uk/projects/fastqc/), followed by a trimming process using trimmomatic v0.38 (PMID: 24695404) to obtain high-quality reads. Qualified reads were then mapped to GRCh38 with STAR v2.7.0e (PMID: 23104886), and subsequently went through an indexing process with samtools v1.10 (PMID: 33590861). Finally, the aligned files were used for generating gene counts by featureCounts programme under subread v2.0.0 (PMID: 24227677). All the mapping process was done on the Hawk high performance computing system based at Cardiff University (https://portal.supercomputing.wales/index.php/about-hawk/).

For downstream analysis, the generated gene counts matrix was filtered, to assess genes expressed in at least 50% of the samples. DESeq2 was next applied for differential expression analysis between groups of comparisons (PMID: 25516281). A Benjamini–Hochberg adjusted p value of $< 0.05$ was considered as statistically significant. To visualize the differential expression results, ggplot2 v3.3.6 package was used to generate volcano plots for the comparisons of interest (https://ggplot2.tidyverse.org). Downstream analysis and visualization were done on R v4.0.3 (https://www.R-project.org).

**Genotyping PCR**

Primer sequences for PCR are provided in **Supplementary sequences**. Genomic DNA from cultured cells was extracted using the NucleoSpin Tissue kit (Macherey-Nagel 740952). Finally, DNA was eluted in 60 µl elution buffer. Genotyping PCR was performed with Q5 PCR DNA polymerase (NEB M0491) plus GC enhancer using 2-4 µl of genomic DNA in 50 µl reaction. PCR products were analyzed on 2-2.5% agarose gels and imaged with a ChemiDoc MD Imaging System (Bio-Rad).

**Western blot**

Cells were washed twice with ice-cold PBS and lysed in RIPA lysis buffer (1% NP-40, 0,1% SDS, 0.5% Sodium Deoxycholate, in 1x TBS) with protease inhibitors (25 µg/ml chymostatin,

25 µg/ml leupeptin, 25 µg/ml antipain hydrochloride, 25 µg/ml pepstatin A). Protein concentration was measured using DC™ Protein Assay Kit I (Bio-Rad, 5000111) and 10 – 15 µg of protein was loaded on Mini-Protein TGX Stain-Free gels (Bio-Rad, 1610181, 1610183, 1610185 and 5678094) and run at 120V in 1x SDS-Page running buffer. After running, gels were activated using the ChemiDoc MD Imaging System (Bio-Rad) and transferred onto 0.45 µm Low Fluorescence PVDF membranes (Bio-Rad, 1704274). Membranes were blocked with 5% skim milk in 0.1% Tween-20 in TBS (TBS-T) for 45 min and subsequently incubated with primary antibodies diluted in blocking buffer overnight at 4ºC. Membranes were washed 3x in TBS-T and incubated with secondary antibodies at room temperature for 45 min. Membranes were washed 3x with TBS-T and incubated with Clarity Western ECL substrate (Bio-Rad, 1705061) and imaged using a ChemiDoc MD Imaging System (Bio-Rad).

**PI4P staining and analysis**

Cellular PI4P staining was performed with mouse anti-PI4P IgM antibody (Echelon, Z-P004). Briefly, cells were fixed with 2% paraformaldehyde (Electron Microscopy Sciences, 15710) in either 1x PBS or culture medium for 15 min at room temperature and permeabilized with 20 µM digitonin (Sigma-Aldrich, 11024-24-1) for 5 min. Cells were then blocked for 1h with a blocking solution containing 1% fatty acid free BSA (Sigma, a-3803) in 1x PBS. Subsequently, cells were incubated with anti-PI4P antibody (1:200), washed 3 times with PBS, then incubated with Alexa FluorTM 488 labeled goat anti-mouse IgM secondary antibody (1:200, ThermoFisher A21042), and washed again at room temperature. Images were taken with Nikon Eclipse Ti-E microscope and analyzed with ImageJ software.

**Lipid droplet staining**

A431 and A549 wild-type and Seipin/BSCL2 degron cells were seeded onto Ibidi 8-well Labtek dishes (Ibidi, 155409) and treated for 24 h with DMSO or pico_cvxIAA. During the final 2 h, 0.2 mM oleic-acid was added as OA:BSA complex. Lipid droplets were stained with the lipid droplet dye LD540 (1:2,000) by adding the dye to medium for the final 20 min. Lipid droplets were imaged by Nikon Eclipse Ti-E widefield microscope using a Plan Apo VC 100x oil DIC N2 objective.

**Generation of H9 AID cells for POGZ**

H9 ES cells were detached as single cells from the culture dishes with StemPro Accutase (Thermo Fisher A1110501) and washed with PBS. Cells were electroporated using the Neon transfection system (Invitrogen). A total of $2.5x10^6$ cells and plasmid mixture, containing Alt-R Cas9 nuclease, sgRNA targeting POGZ (TCTGATGGAGATTTGAGTGT TGG), Electroporation enhancer, and the donor template plasmid (POGZ-miniIAA7-GFP), were electroporated in a 100 uL tip with 1100 V, 20 ms, and 2x pulse settings. Electroporated H9 ES cells were plated on Matrigel coated 35mm dishes in mTeSR medium containing 10 µM ROCK inhibitor Y-27632 2HCL (Selleckchem S1049) and 1 µM Alt-R HDR Enhancer. After 24 h, medium was changed to mTeSR medium without ROCK inhibitor or HDR enhancer, and the cells were further cultured until 72 h. The HDR efficiency was checked with FACS analysis. GFP+ cells were single-cell sorted into 96 well plate, and clones were expanded and checked with gPCR for identifying homozygous or heterozygous tagging. One homozygously tagged clone was selected to introduce *At*AFB2 (F74A)-SNAPf-weakNLS with BSD selection marker into *AAVS1* locus by electroporation, followed by blasticidin selection for 2 weeks.

**H9 cells culture and one-step generation of AID clones**

H9 cells (WiCell, WIC-WA09-RB-001) were cultured in mTeSR Plus culture medium (Stem cell technologies, 100-0276) on plates coated with Matrigel (Corning, 356231) diluted 1:200 in DMEM/F-12 (Gibco, 31331-028). Cells were detached by washing carefully 1-2x with PBS (Corning, 21-040-cv), and subsequently treated with 0.5 µM EDTA (Invitrogen 15575020, diluted 1:1,000 in PBS) for 4-5 min at RT. Alternatively, cells were passaged with StemPro EZPassage Disposable Stem Cell Passaging Tool (Invitrogen, 23181-010) and scraping in mTeSR Plus medium.

For transfection of H9 cells, the medium of H9 cells was changed to fresh mTeSR Plus medium 3-5 h before passaging. H9 cells were washed with PBS, incubated with StemPro Accutase (Thermo Fisher, A1110501) for 5-6 min at 37ºC, and centrifuged for 4 min at 1,000 rpm. The cell pellet was resuspended in 2 ml mTeSR Plus medium with 10 µM Y27632 (Selleckchem, S1049). Cells were counted using a TC10 Automated cell counter (Bio-Rad) and histograms were checked to avoid counts with abnormal distributions. $3x10^5$ - 5 $x10^5$ cells were passaged to a 6-well in mTeSR medium with 10 µM Y27632. On day 1, medium was changed to 2 ml of fresh mTeSR Plus medium with 10 µM Y27632. H9 cells were transfected using 5 µl Lipofectamine STEM transfection reagent (Thermo Fisher, STEM00015) with 2.5 µg plasmids. At 4-6 h post-transfection, medium was changed to fresh medium with 10 µM Y27632 and different concentrations of M3814 (0, 0.25 and 1 µM). On day 2, medium with 10 µM Y27632, 0.5 µg/ml puromycin and indicated concentration of M3814 was added to the cells. Medium was changed to mTeSR Plus medium supplemented with M3814 and puromycin on day 3. On day 4 puromycin medium was added to the cells, followed by 2-3 days of 10 µg/ml Blasticidin selection until colonies were picked. Before picking colonies, medium was changed to fresh mTeSR Plus medium without antibiotics and 6-8 colonies per transfection were picked and transferred to a 4-well dish with 0.5 ml of mTeSR Plus medium. Medium of colonies was changed every 2 days for clonal growth.

**H9 cells for LD staining**

H9 WT and Seipin/BSCL2 degron cells were seeded in mTeSR with Y27632 onto Matrigel coated Ibidi 8-well Labtek dishes (Ibidi, 155409) and treated for 24 h with DMSO or pico_cvxIAA. During the final 4h 0.4 mM oleic acid (prepared as a 1 mM OA-BSA complex at a 8:1 molar ratio to BSA in serum free DMEM) was added to the cells. Lipid droplets were stained with the lipid droplet dye LD540 by adding the dye to mTeSR medium for the final 20 min (1:2000, Princeton BioMolecular Research). Lipid droplets were imaged using a Nikon Eclipse Ti-E widefield microscope with a Plan Apo VC 100x oil DIC N2 objective. Z-stacks were taken with a 0.3 µm interval. Images represent maximal projections and brightness and contrast were adjusted in ImageJ.

**Differentiation of H9 cells to neurons**

Human neurons were derived by differentiating human H9 ES cells using a small molecule cocktail as described before, with minor adjustments.[37] N2B27 medium (N2B27 basal medium; 50% DMEM/F12 and 50% Neurobasal medium supplemented with 0.5x N2 and 0.5x B27, 1 mM GlutaMAX, and 1x Penicillin-Streptomycin) was used throughout the differentiation protocol. Before induction, H9 ES cells were passaged in mTeSR™ Plus and replated to form a uniform monolayer of cells. When the cells had reached around 95% confluence, medium was replaced with dual SMAD inhibition medium: N2B27 supplemented with 2 µM dorsomorphin (Selleckchem S7306) and 10 µM SB431542 (Sigma S4317). On day 10, the cells were replated in 1:2 ratio using 200 U/mL Collagenase IV (Gibco) onto Matrigel-coated dishes in N2B27 supplemented with 10 µM Y-27632 (Selleckchem S1049). From day 11 to day 20, N2B27 was supplemented with 100 ng/mL of FGF8 (PeproTech AF-100-25). On day 20, cells were detached using 0.5 mM EDTA and replated in N2B27 at 1:8 ratio. The next day following the split, N2B27 was supplemented with 20 µM DAPT (Selleckchem S2215) and the medium was replaced every 2 days.

**Preparing neuronal samples for WB, PMP70 staining and live-cell imaging**

The H9 ES cell-derived neurons were collected on day 25 of differentiation. For WB, neurons were grown in 35mm dishes. Medium was removed, and neurons were washed once with 1mL DMEM/F12 and once with 1mL PBS. Neurons were detached with cell scraper, collected into a 1.5mL Eppendorf tube, and centrifuged at 300xg for 3min. The supernatant was removed, and cells were lyzed in RIPA buffer for WB analysis. For PMP70 staining, neurons were grown on Matrigel-coated coverslip. Medium was removed, and neurons were washed once with DMEM/F12 and once with sterile PBS. Neurons were fixed with 2% PFA for 15 min. For live-cell imaging, neurons were grown on ibidi plate (Cat.80826) and processed for imaging.

**PMP70 immunofluorescence microscopy**

Fixed cells were quenched with 50 mM $NH_4Cl$ for 10 min, permeabilized with 0.1% saponin (Sigma S4521) in PBS for 10 min, and blocked by incubation with 10% FBS in PBS for 30 min. The cells were then stained with anti-PMP70 antibodies (Sigma, SAB4200181) for 1 h and Alexa Fluor 568-conjugated secondary antibodies (Thermo Fisher, A11004) for 30 min. Prior to each antibody incubation, the cells were washed with 0.1% saponin in PBS. Cells mounted with Mowiol/DABCO (Calbiochem 475904/Sigma D2522) were imaged with a confocal Leica Stellaris 8 inverted microscope using 63x HC PL APO CS2 oil objective NA 1.40.

**Statistics and reproducibility**

Graphpad Prism 9 (Graphpad Software, Inc.) was used to generate graphs. Quantitative data are presented as mean ± S.D. Results were validated in at least two cell lines for each endogenous target.

### Supplementary notes

#### Suppl. Note 1: Comparison of different AID components.

Several pitfalls have been described regarding the AID technique, including basal degradation before induction, inefficient inducible degradation and high inducer concentration required (500 µM IAA). Different possibilities for improving AID components were recently published.[8,9,13] Systematic comparisons of the available options were conducted here.

We first compared the different auxin receptors with Seipin-miniIAA7-mEGFP as the substrate **(Suppl. Fig. 1a).** As reported, the auxin receptor *Os*TIR1 caused severe basal degradation and this could be avoided by using *At*AFB2 or *Os*TIR1(F74G) mutant **(Suppl. Fig. 1b)**.[8,9] The new *At*AFB2(F74A) mutant, like *Os*TIR1(F74G), showed no basal degradation and high sensitivity to the engineered inducers cvx_IAA and pico_cvxIAA **(Suppl. Fig. 1b-g)**. The F74G or F74A mutation lost binding to IAA.[11] We thus envisage that basal degradation of *Os*TIR1 might be caused by an unknown endogenous ligand analogous to IAA with low activity, and *Os*TIR1(F74G) with low affinity for IAA would thus loose the binding to this unknown ligand to reverse the basal degradation.[11] The enhanced sensitivity of engineered inducers might result from their higher binding affinity[12] and improved membrane-permeability compared to IAA that has poor membrane permeability at natural pH[51].

*Os*TIR1(F74G) behaved as *At*AFB2(F74A) in the test, while *Os*TIR1(F74A) showed residual amount of basal degradation but had a 10-fold higher sensitivity to pico_cvxIAA (**Suppl. Fig. 1b-g**). *Os*TIR1(F74A) might thus be beneficial when ligand concentration is limiting, such as in certain *in vivo* applications.

Rapid inducible degradation with AID can be reversed by simply washing out the inducers. *At*AFB2(F74A) treated with 5 µM cvx_IAA, and *At*AFB2 with 500 µM IAA, showed the best reversibility **(Suppl. Fig. 1h-i)**. Pico_cvxIAA showed poor reversibility, likely due to the retention of residual inducer in the cells after washout and the high sensitivity of the auxin receptors to pico_cvxIAA **(Suppl. Fig. 1h-i)**.

A small tag likely has less steric hinderance and reduced impact on the target protein functions. Interestingly, the small degron tag miniIAA7-3xFlag showed minimal impact on EGFP stability before induction and the fastest inducible degradation in comparison to miniAID, miniIAA7, and other small miniIAA7 fusion tags tested **(Suppl. Fig. 1k-l)**. These results emphasize that the choice of degron tag may have a major impact on both the target protein stability and its rapid inducible degradation. In addition, simple comparisons of different degron tagging technologies without proper optimization could lead to biased conclusions.[52] We also noticed that the miniIAA7 degron was incorrectly used in the previous literature, resulting in an erroneous conclusion.[9]

Pico_cvxIAA was the most sensitive inducer and showed negligible off-target activity at its effective concentration (0.5 µM) compared to cvxIAA (5 µM) and IAA (500 µM) **(Suppl. Fig. 1m-n)**. We thus used 0.5 µM pico_cvxIAA as the inducer of proteolysis for all the subsequent experiments. With IAA, FABP4 was upregulated by more than 8-fold after 24 h treatment, which was not observed with the other inducers **(Suppl. Fig. 1m-n)**. It should be noticed that the lower concentration of the inducer is one of the main reasons for lower off-target effects, as cxv_IAA and pico_cvxIAA, but not IAA, at concentration above 50 µM showed clear cell toxicity (data not shown).

**Suppl. Note 2: Optimization of coIN and HDR enhancers.**

Simultaneous introduction of two independent genomic modifications results in high coincidence in the same cells, provided that they are mediated by a similar DNA repair pathway (HDR or NHEJ)[26,27] CoIN, or co-selection, uses this phenomenon to effectively enrich cells with both genomic modifications. The mechanism is harnessed here for one-step introduction of two AID components. Degron tagging was mediated by HDR. An HDR-mediated *AAVS1* safe harbor integration system (2 plasmids) was thus chosen to overexpress the auxin receptor *At*AFB2(F74A)-mCherry (**Suppl. Fig. 2a)**. After selection with puromycin, the majority of cells expressed *At*AFB2(F79A)-mCherry (>99%) and integration was effectively mediated by HDR (sgAAVS1) instead of random integration (sgCtrl) or homology-independent targeted insertion (sgSec61B)[49], especially in the presence of HDR enhancers **(Suppl. Fig. 2b)**. For degron-tagging through coIN, an endogenous degron-GFP tagging pair (2 plasmids) was co-transfected with the *AAVS1* integration system **(Suppl. Fig. 2c-d)**. *At*AFB2(F74A)-mCherry expressing cells were selected with puromycin and the efficiency of degron-GFP tagging in the selected cells was measured by FACS. The efficiency of endogenous tagging increased with a higher amount of endogenous tagging plasmids. Tagging efficiency reached a plateau at 1:3 ratio and a further increase of the ratio decreased the number of puromycin resistant cells **(Suppl. Fig. 2d-e)**. The ratio of 1:3 was thus chosen for effective degron-tagging with a high yield of puromycin selected cells. *At*AFB2(F74A)-mCherry expression was commonly close to 100% and will not be described separately below.

Inhibitors targeting DNA-dependent protein kinase (DNA-PK) and 53BP1 were used as they have been consistently reported to increase HDR efficiencies. Several inhibitors with different properties were tested to improve degron-tagging efficiency in coIN. These included M3814 (a small-molecule inhibitor of DNA-PK)[17], i53 (a small-peptide inhibitor of 53BP1)[18], and Cas9-53BP1-DN1S (a Cas9 fusion protein that was designed to inhibit 53BP1 locally at Cas9 cut sites)[19]. In addition, XL413 (a CDC7 inhibitor controlling the cell cycle) was included as it was recently identified in a large screen to improve HDR[20]. Two of the inhibitors, M3814 and i53, effectively increased degron-GFP tagging efficiencies in coIN (7.5-fold increase in GFP levels with 1 µM M3814 and 2.6-fold with i53 overexpression) **(Suppl. Fig. 3a)**.

I53, but not 1 µM M3814, improved degron-GFP tagging efficiency without reducing cell counts **(Suppl. Fig. 3a)**. Moreover, i53 reduced the concentration of M3814 needed from 1 µM to 0.25 µM to achieve high tagging efficiency (**Suppl. Fig. 3b)**. 1 µM M3814 and i53 plus 0.25 µM M3814 were thus chosen for further tests.

**Suppl. Note 3: Comparison of Sh_ble and BSD as S2.**

*Streptoalloteichus hindustanus* bleomycin (Sh_ble) and Blasticidin S deaminase (BSD) genes, that confer resistance to Zeocin and Blasticidin respectively, were initially selected as S2 due to their small size (369 bp and 396 bp) **(Suppl. Fig. 4a** and **Fig. 2f)**. The P2A-S2 cassettes do not have a promoter and thus will not likely be expressed through random integration. In cells with degron tagging, the selection marker is transcribed under the control of an endogenous promoter and is translated at 1:1 ratio with the endogenous target through P2A self-cleavage.[53] Single clones were initially isolated as in **Fig. 1c** without HDR enhancer. Using Sh_ble as S2, homozygous clones were effectively enriched for 2 targets (SAC1 and DHC1) **(Suppl. Fig. 4b)**. However, further tests showed that low-expressing BSCL2/Seipin clones died out and high-expressing ones, such as SEC61B, LMNA and MYH9,[8] had a high number of heterozygous clones **(Suppl. Fig. 4b)**.[8] Moreover, SAC1 and DHC1 clones grew more slowly than the high-expressing clones during Zeocin selection with clear indications of cell stress (data not shown). Further tests show that HDR enhancer M3814 (1 µM) substantially improved the efficiency of homozygous tagging for the high-expressing targets with Sh_ble as S2 **(Suppl. Fig. 4b)**. These results, together with results using BSD as S2 **(Fig. 2i)**, consistently show that HDR enhancers substantially improved homozygous degron tagging efficiencies in one-step generation of AID cells.

BSD catalyzes the degradation of Blasticidin[53] while Sh_ble binds stoichiometrically to Zeocin[54]. The catalytic feature of BSD might explain its higher sensitivity to enrich the low-expressing Seipin/BSCL2 **(Fig. 2i)**. The choice of S2 in the experiment would be a consideration of drug sensitivity in the cell line combined with the expression level of the target protein.

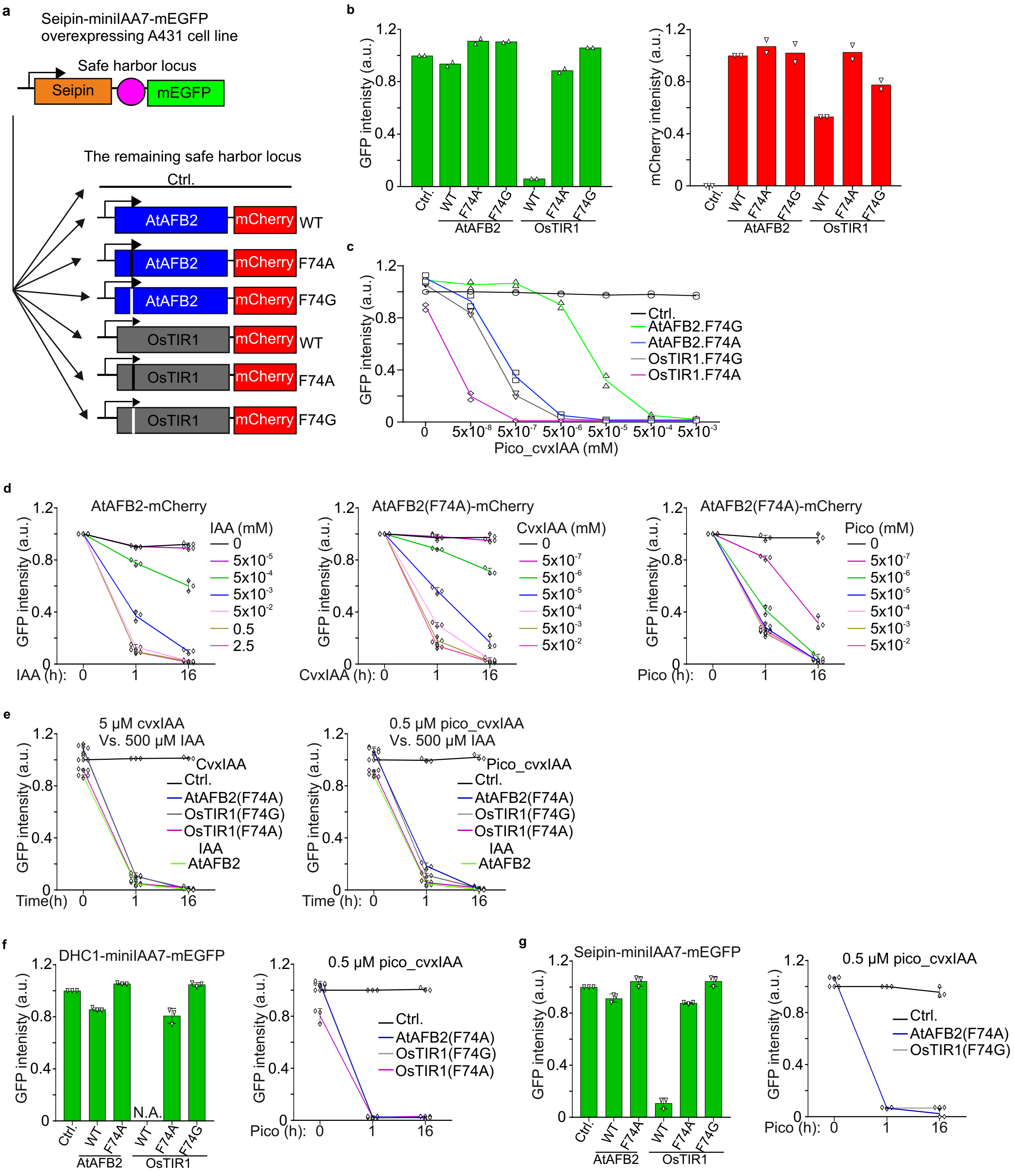

a
Seipin-miniIAA7-mEGFP overexpressing A431 cell line
Safe harbor locus
Seipin
mEGFP
The remaining safe harbor locus
Ctrl.
AtAFB2
mCherry
WT
F74A
F74G
OsTIR1
b
GFP intenisty (a.u.)
mCherry intenisty (a.u.)
c
Pico_cvxIAA (mM)
Ctrl.
AtAFB2.F74G
AtAFB2.F74A
OsTIR1.F74G
OsTIR1.F74A
d
AtAFB2-mCherry
AtAFB2(F74A)-mCherry
IAA (mM)
CvxIAA (mM)
Pico (mM)
IAA (h):
CvxIAA (h):
Pico (h):
e
5 μM cvxIAA
Vs. 500 μM IAA
0.5 μM pico_cvxIAA
Vs. 500 μM IAA
CvxIAA
Pico_cvxIAA
AtAFB2(F74A)
OsTIR1(F74G)
OsTIR1(F74A)
IAA
AtAFB2
Time(h)
Time (h):
f
DHC1-miniIAA7-mEGFP
N.A.
g
Seipin-miniIAA7-mEGFP

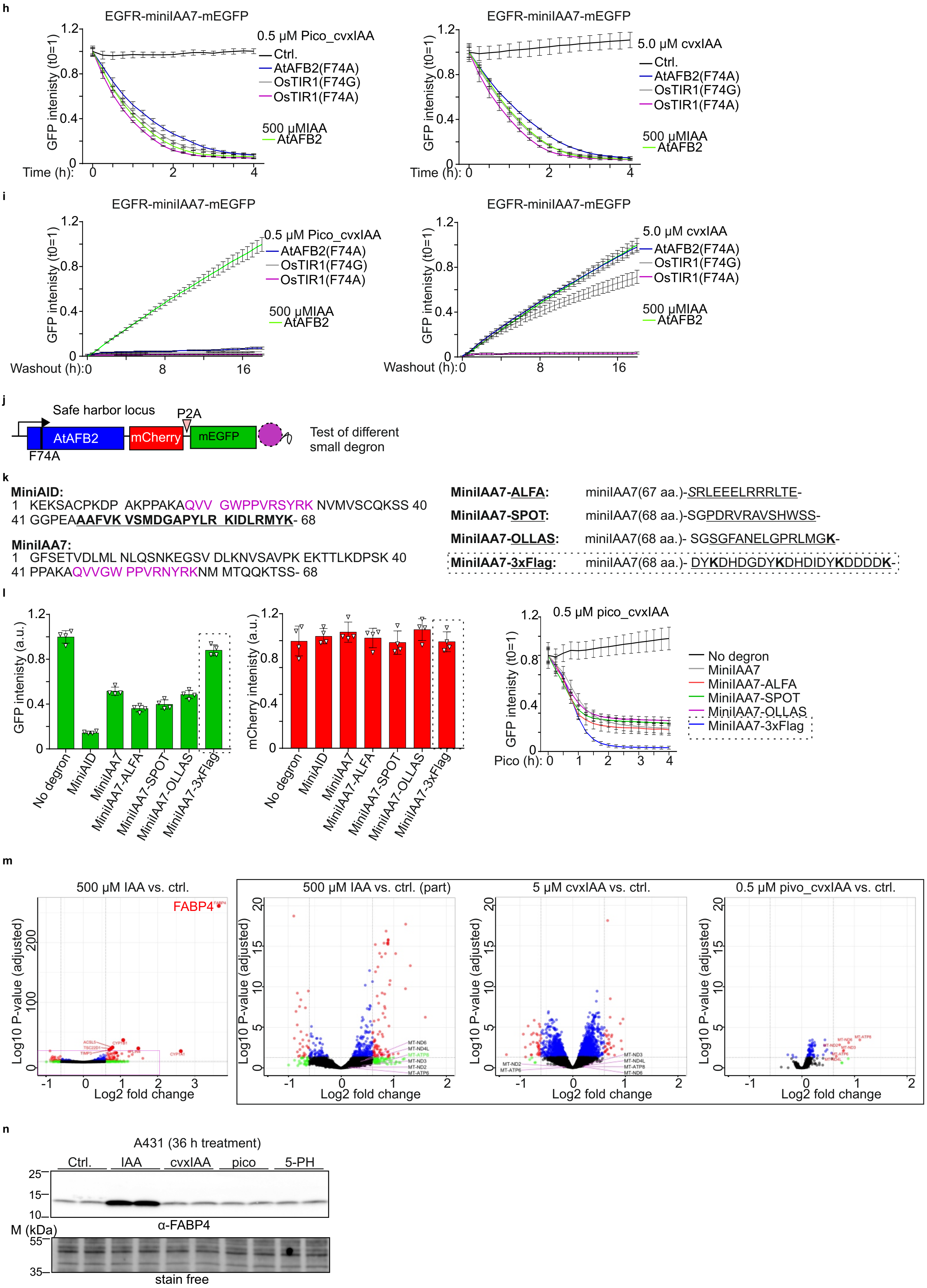
h
EGFR-miniIAA7-mEGFP
0.5 µM Pico_cvxIAA
Ctrl.
AtAFB2(F74A)
OsTIR1(F74G)
OsTIR1(F74A)
500 µMIAA
AtAFB2
GFP intenisty (t0=1)
Time (h):
EGFR-miniIAA7-mEGFP
5.0 µM cvxIAA
Ctrl.
AtAFB2(F74A)
OsTIR1(F74G)
OsTIR1(F74A)
500 µMIAA
AtAFB2
GFP intenisty (t0=1)
Time (h):
i
EGFR-miniIAA7-mEGFP
0.5 µM Pico_cvxIAA
AtAFB2(F74A)
OsTIR1(F74G)
OsTIR1(F74A)
500 µMIAA
AtAFB2
GFP intenisty (t0=1)
Washout (h):
EGFR-miniIAA7-mEGFP
5.0 µM cvxIAA
AtAFB2(F74A)
OsTIR1(F74G)
OsTIR1(F74A)
500 µMIAA
AtAFB2
GFP intenisty (t0=1)
Washout (h):
j
Safe harbor locus
P2A
AtAFB2
F74A
mCherry
mEGFP
Test of different
small degron
k
MiniAID:
1 KEKSACPKDP AKPPAKAQVV GWPPVRSYRK NVMVSCQKSS 40
41 GGPEAAAFVK VSMDGAPYLR KIDLRMYK- 68
MiniIAA7:
1 GFSETVDLML NLQSNKEGSV DLKNVSAVPK EKTTLKDPSK 40
41 PPAKAQVVGW PPVRNYRKNM MTQQKTSS- 68
MiniIAA7-ALFA: miniIAA7(67 aa.)-SRLEEELRRRLTE-
MiniIAA7-SPOT: miniIAA7(68 aa.)-SGPDRVRAVSHWSS-
MiniIAA7-OLLAS: miniIAA7(68 aa.)- SGSGFANELGPRLMGK-
MiniIAA7-3xFlag: miniIAA7(68 aa.)- DYKDHDGDYKDHDIDYKDDDDK-
l
GFP intenisty (a.u.)
No degron
MiniAID
MiniIAA7
MiniIAA7-ALFA
MiniIAA7-SPOT
MiniIAA7-OLLAS
MiniIAA7-3xFlag
mCherry intenisty (a.u.)
0.5 µM pico_cvxIAA
GFP intenisty (t0=1)
Pico (h):
No degron
MiniIAA7
MiniIAA7-ALFA
MiniIAA7-SPOT
MiniIAA7-OLLAS
MiniIAA7-3xFlag
m
500 µM IAA vs. ctrl.
FABP4
Log10 P-value (adjusted)
Log2 fold change
500 µM IAA vs. ctrl. (part)
5 µM cvxIAA vs. ctrl.
0.5 µM pivo_cvxIAA vs. ctrl.
n
A431 (36 h treatment)
Ctrl.
IAA
cvxIAA
pico
5-PH
α-FABP4
M (kDa)
stain free

Suppl. Fig.1

**Suppl. Fig. 1: Comparison of AID components.**
**a,** scheme of the genomic modifications in cells to compare different auxin receptors using overexpressed Seipin-miniIAA7-mEGFP as substrate.
**b,** basal levels of Seipin-miniIAA7-mEGFP and expression levels of mCherry tagged auxin receptors before induction, analyzed by FACS. N=2 independent experiments.
**c,** levels of Seipin-miniIAA7-mEGFP after 16 h treatment of pico_cvxIAA at indicated concentrations in cells expressing different auxin receptors, analyzed by FACS. N=2 independent experiments.
**d,** levels of Seipin-miniIAA7-mEGFP after treatment with IAA (left) or engineered IAA (middle and right) for 0, 1 and 16 h at indicated concentrations, analyzed by FACS. N=3 independent experiments.
**e,** levels of Seipin-miniIAA7-mEGFP after treatment with 500 µM IAA (left and right), 5 µM cvxIAA (left) or 0.5 µM pico_cvxIAA (right) for 0, 1 and 16 h in cells expressing different auxin receptors, analyzed by FACS. N=3 independent experiments.
**f,** basal levels (left) and inducible degradation (right) of endogenous DHC1-miniIAA7-mEGFP in cells expressing different auxin receptors, analyzed by FACS. N=3 independent experiments.
**g,** basal levels (left) and inducible degradation (right) of endogenous Seipin-miniIAA7-mEGFP in cells expressing different auxin receptors, analyzed by FACS. N=3 independent experiments.
**h,** rapid inducible degradation of endogenous EGFR-miniIAA7-mEGFP (heterozygous tagging) after treatment with 500 µM IAA (left and right), 5 µM cvxIAA (left) or 0.5 µM pico_cvxIAA (right) in cells expressing different auxin receptors, analyzed by live-cell imaging. N=4 fields. Representative of 2 independent experiments.
**i,** recovery from washout of endogenous EGFR-miniIAA7-mEGFP (heterozygous tagging) in cells after 16 h treatment with 500 µM IAA (left and right), 5 µM cvxIAA (left) or 0.5 µM pico_cvxIAA (right). N=4 fields. Representative of 2 independent experiments.
**j,** scheme of genetic modifications to compare different small degron tags. P2A: self-cleavage peptide.
**k,** amino acid sequences of different small degrons. Magenta: core sequences for the auxin receptor binding; bold and underlined: sequence derived from the dimerization domain in miniAID; framed: miniIAA7-3xFlag used to establish AID clones.
**l,** basal levels of mEGFP fusions (left), expression levels of *At*AFB2(F74A)-mCherry (middle), and the inducible degradation of mEGFP fusions (right), analyzed by live-cell imaging. N=4 fields. Representative of 2 independent experiments.
**m,** volcano plots of RNAseq in A431 wild-type cells treated with different inducers for 24 h compared to control. Full plot of IAA treated cells is shown on the left with 7 targets out of the range for comparison (magenta frame); FABP4 highlighted for WB analysis. No target is out of this range for cells treated with the other 2 inducers. Threshold for different colors: fold change at >1.5 or <0.5, P value at 0.05.
**n,** WB analysis of FABP4 in A431 cells treated with different inducers for 36 h. 500 µM IAA, 5.0 µM cvxIAA, 0.5 µM pico_cvxIAA or 1 µM 5-PH was used.

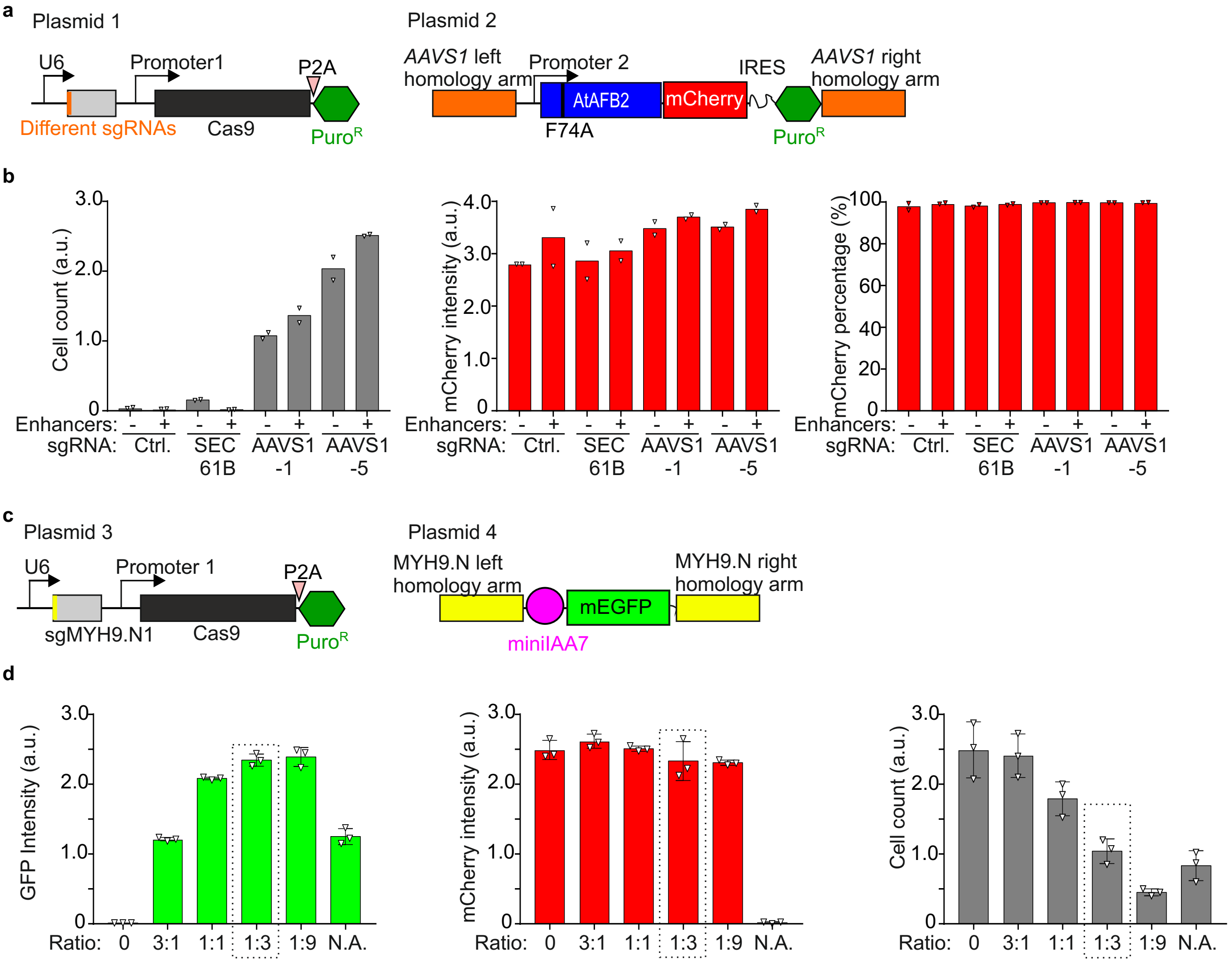


**e**

| |
|---|
| ***AAVS1*:MYH9 Ratio=1:3** |
| (for a 12-well, total of 1 µg plasmids) |
| Plasmid 1 (µg): 0.1 |
| Plasmid 2 (µg): 0.15 |
| Plasmid 3 (µg): 0.3 |
| Plasmid 4 (µg): 0.45 |

**Suppl. Fig. 2: Optimization of coIN for one-step generation of AID cells.**

**a,** scheme of the plasmid pair used in (b) to test the HDR-mediated *AAVS1* integration efficiency.

**b,** cell count (left), mCherry intensity (middle) and percentage of mCherry positive cells (right) in cell pools generated with indicated sgRNAs. Cells were transfected with Plasmid 1 and plasmid 2 at 2:3 ratio (µg: µg). Enhancers: i53 plus 0.25 µM M3814 as HDR enhancers (see Suppl. Fig. 3 for a description of HDR enhancers).

**c,** scheme of the plasmid pair used in (d) to test HDR-mediated endogenous miniIAA7-mEGFP tagging efficiency of MYH9.

**d,** levels of GFP (MYH9 tagging efficiency), mCherry (*At*AFB2(F74A) expression level), and cell counts (*AAVS1* integration efficiency) in cell pools generated with indicated ratio of *AAVS1*: MYH9 tagging plasmid pairs (plasmid 1+2: plasmid 3+4). Wildtype A431 cells were co-transfected with the 4 plasmids and selected for stable puromycin-resistant cell pools before analysis (except for group N.A.). N.A. no *AAVS1* integration plasmid pair.

**e,** amount of 4 plasmids used at 1:3 ratio (*AAVS1*: endogenous loci) for transfection of a 12-well of cells.

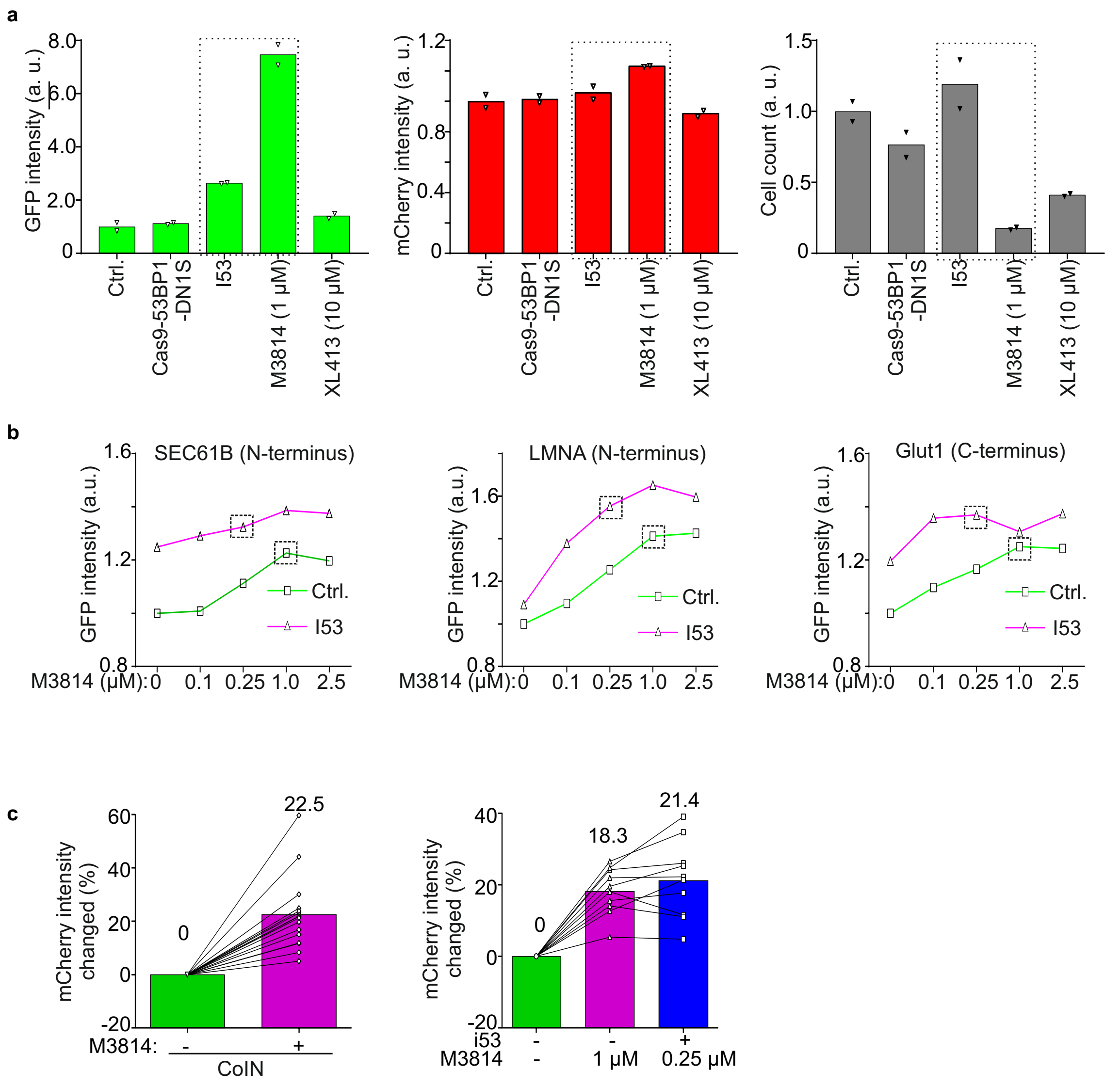


**Suppl. Fig. 3: Test of different HDR enhancers in coIN.**

**a,** levels of GFP (MYH9 tagging efficiency), mCherry (AtAFB2(F74A) expression level), and cell count (cell toxicity) in stable cell pools using different HDR enhancers. Stable cell pools were generated by coIN with *AAVS1*:MYH9 at 1:3 ratio. N=2 technical repeats. Frames indicate M3814 and i53 used for further analysis.

**b,** endogenous tagging efficiencies of SEC61B (left), LMNA (middle) and Glut1 (right) in cell pools generated by coIN with or without i53 expression plus different concentration of M3814. Frames indicate 1 μM M3814 and i53 plus 0.25 μM M3814 chosen for later experiments.

**c,** mCherry expression levels with indicated HDR enhancers. Results from the same experiment as **Fig. 2 c** and **e.** N=16 (left) and 10 (right). Numbers above columns indicate mean values; lines link the same endogenous tagging pairs.

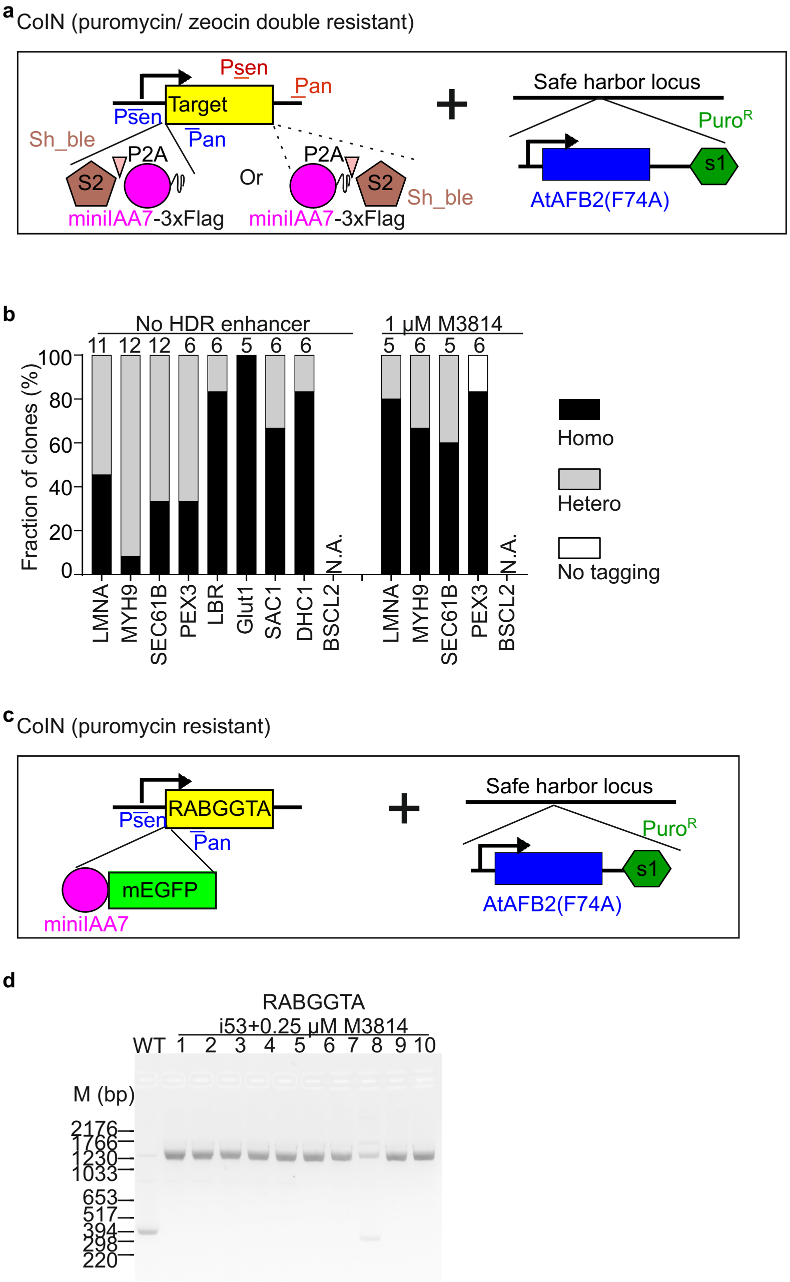


**Suppl. Fig. 4: One-step generation of AID clones in A431 cells with Sh_ble as S2 or without S2.**

**a,** scheme of the genomic modifications in one-step generation of AID cells using Sh_ble as S2.

**b,** statistics of genotyping PCR results for 9 targets without HDR enhancer, and 5 targets with 1 μM M3814 as the HDR enhancer. N.A. not available due to cell death after Zeocin (S2) selection. Numbers indicate total amount of clones analyzed.

**c,** scheme of the genomic modifications in one-step generation of AID cells targeting RABGGTA without S2.

**d,** genotyping PCR results for RABGGTA clones generated without S2. WT: wild-type

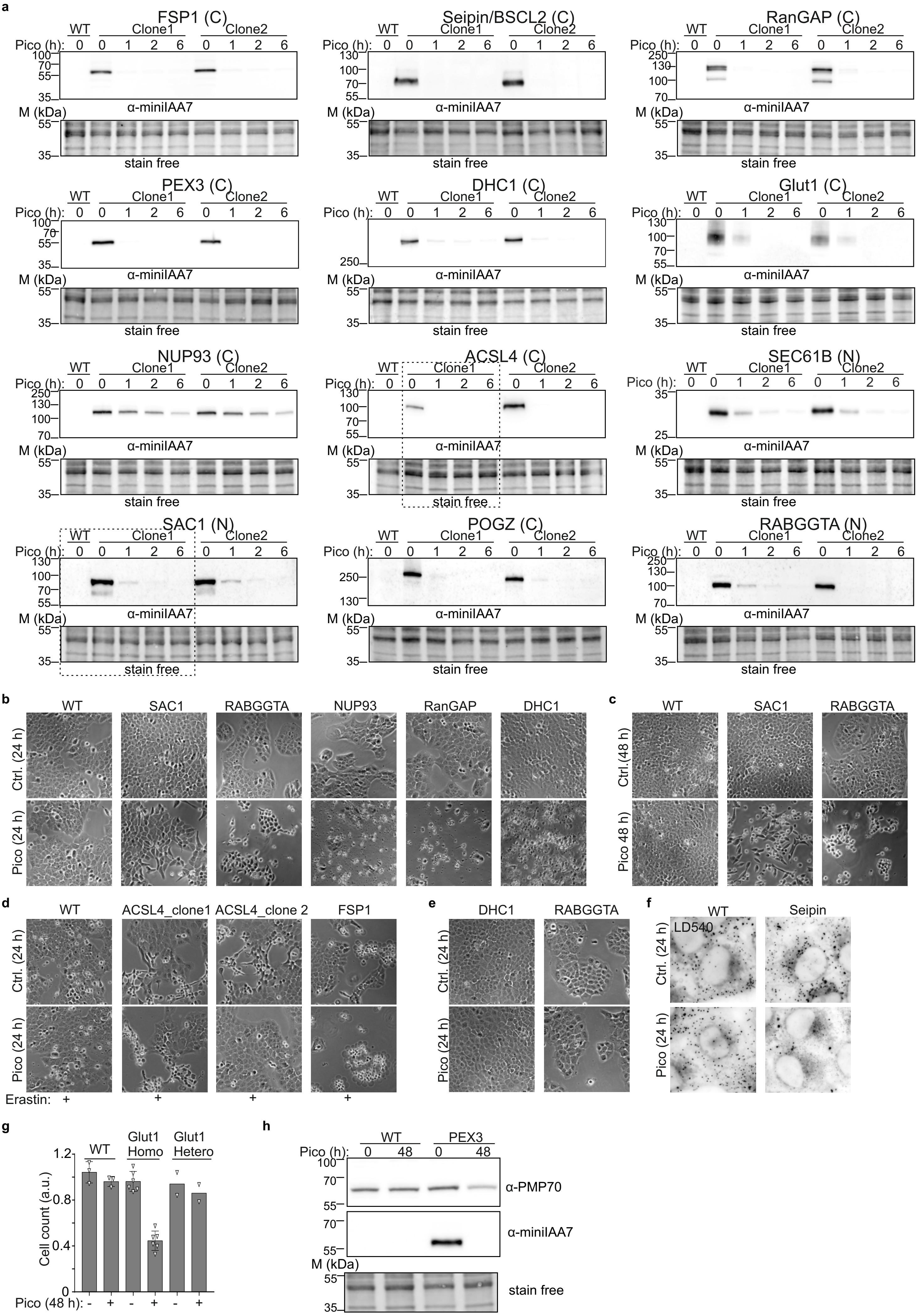
a
FSP1 (C)
Seipin/BSCL2 (C)
RanGAP (C)
PEX3 (C)
DHC1 (C)
Glut1 (C)
NUP93 (C)
ACSL4 (C)
SEC61B (N)
SAC1 (N)
POGZ (C)
RABGGTA (N)
WT
Clone1
Clone2
Pico (h): 0 0 1 2 6 0 1 2 6
M (kDa)
α-miniIAA7
stain free
b
WT
SAC1
RABGGTA
NUP93
RanGAP
DHC1
Ctrl. (24 h)
Pico (24 h)
c
WT
SAC1
RABGGTA
Ctrl.(48 h)
Pico 48 h)
d
WT
ACSL4_clone1
ACSL4_clone 2
FSP1
Ctrl. (24 h)
Pico (24 h)
Erastin: + + + +
e
DHC1
RABGGTA
Ctrl. (24 h)
Pico (24 h)
f
WT
Seipin
LD540
Ctrl. (24 h)
Pico (24 h)
g
WT
Glut1 Homo
Glut1 Hetero
Cell count (a.u.)
1.2
0.8
0.4
0
Pico (48 h): - + - + - +
h
WT
PEX3
Pico (h): 0 48 0 48
α-PMP70
α-miniIAA7
M (kDa)
stain free

Suppl. Fig.5

**Suppl. Fig. 5: WB and functional analysis of the A431 degron cell lines generated with HDR enhancers.**
**a,** WB analysis of inducible degradation with anti-miniIAA7 antibody. 2 homozygous clones identified in gPCR generated with HDR enhancers for each target. Frames highlight the results of SAC1 clone 1 shown in Fig. 2j**,** and ASCL4_clone 1 that might be heterozygously tagged with no clear phenotypic change after induction in (d). N and C in brackets indicate terminus of degron-tagging.
**b-c,** representative live-cell images showing morphological changes and cell death in homozygous clones after 24 h (b) and 48 h (c) of induction. N=8 (SAC1), 9 (RABGGTA), 6 (NUP93), 5 (DHC1) and 4 (RANGAP1) clones.
**d,** representative live-cell images showing erastin-induced ferroptosis in homozygous clones after 24 h of induction. FSP1 inhibits while ASCL4 promotes ferroptosis. ASCL4_clone 1 with lower degron-tagged protein level in (a) show no clear phenotypic change. N=2 clones for FSP1.
**e,** live-cell images of heterozygous clones identified in gPCR showing no clear phenotypic changes after 24 h of induction. N=3 (DHC1) and 1 (RABGGTA) clone.
**f,** live-cell images showing lipid droplets stained by LD540 with lots of small and a few big lipid droplets in BSCL2/seipin degron cells after 24 h induction; 0.2 mM of oleic aicd was added during the final 2 h to induce LD formation. N=6 clones.
**g,** growth of cells analyzed by cell counting after 48 h of induction. Homo: homozygous; Hetero: heterozygous; each data point of a Glut1 column representing one clone. N=6 (Homo) and 2 (Hetero) clones.
**h,** WB analysis of peroxisomal membrane protein PMP70 depletion in PEX3 homozygous clones after 48 of induction.
WT: wild-type; pico: 0.5 µM pico_cvxIAA treatment; a.u.: arbitrary unit.

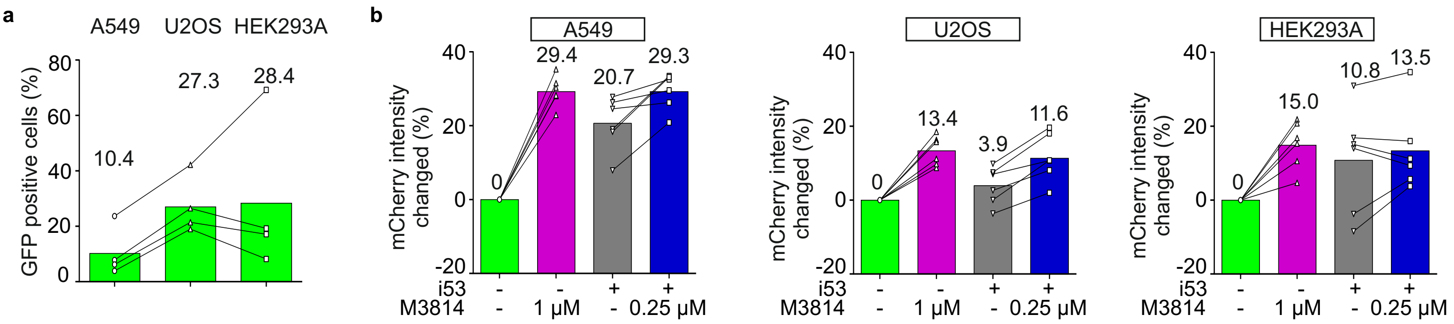


**Suppl. Fig. 6: Evaluation of coIN and HDR enhancers in other human cancer cell lines. a.** percentage of degron-GFP tagging in A549, U2OS and HEK293A cells through conventional tagging without *AAVS1* integration analyzed by FACS. Numbers above columns indicate mean value. Lines link the same endogenous tagging pair. N=4 (2 target proteins with 2 sgRNAs each).

**b,** mCherry expression levels with indicated HDR enhancers in A549, U2OS and HEK293A cells. Results from the same experiment as **Fig. 3 b, d, f.** Numbers above columns indicate mean values; lines link the same endogenous tagging pairs. N=6.

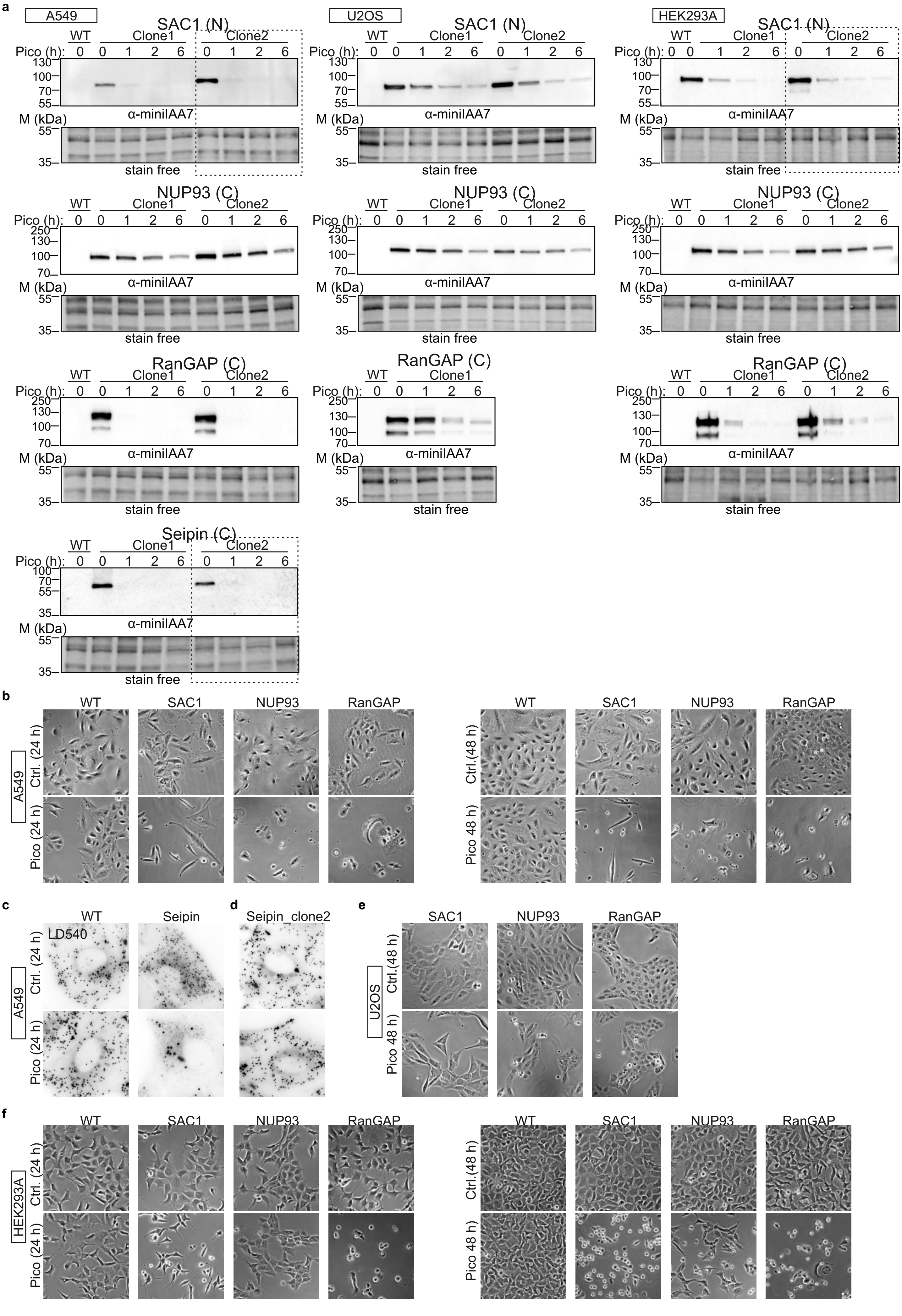
a
A549
U2OS
HEK293A
SAC1 (N)
NUP93 (C)
RanGAP (C)
Seipin (C)
WT
Clone1
Clone2
Pico (h): 0 0 1 2 6 0 1 2 6
M (kDa)
250
130
100
70
55
35
α-miniIAA7
stain free
b
WT
SAC1
NUP93
RanGAP
Ctrl. (24 h)
Pico (24 h)
Ctrl.(48 h)
Pico 48 h)
c
Seipin
LD540
d
Seipin_clone2
e
f

Suppl. Fig.7

**Suppl. Fig. 7: WB and functional analysis of A549, U2OS and HEK293A degron cell lines generated with HDR enhancers.**
**a,** WB analysis of inducible degradation with anti-miniIAA7 antibody. Homozygous clones identified in gPCR generated with HDR enhancers. Frames highlight the results of SAC1 shown in Fig. 3h, and Seipin_clone 2 that might be heterozygously tagged with no clear phenotypic change after induction in (c). N and C in brackets indicate terminus of degron-tagging.
**b,** live-cell images showing morphological and cell-density changes in homozygous clones after 24 h (left) and 48 h (c) of induction in A549 cells. N=7 (NUP93), 5 (both SAC1 and RANGAP1) clones.
**c-d,** live-cell images of lipid droplets stained by LD540 showing lots of small with a few big lipid droplets in A549 BSCL2/seipin degron cells after 24 h pico_cvxIAA treatment (c) and Seipin_clone 2 without clear change (d). 0.2 mM of oleic aicd was added during the final 2 h to induce LD formation. WB of Seipin_clone 2 shown in (a). N=5 (c) and 1(d) clone.
**e,** live-cell imaging analysis of morphological changes in homozygous clones after 48 h of induction in U2OS cells.
**f,** live-cell images showing morphological and cell density changes in homozygous clones after 24 h (left) and 48 h (c) of induction in HEK293A cells. N= 4 (SAC1), 1 (NUP93) and 2 (RANGAP1) clones.
WT: wild-type; pico: 0.5 µM pico_cvxIAA treatment.

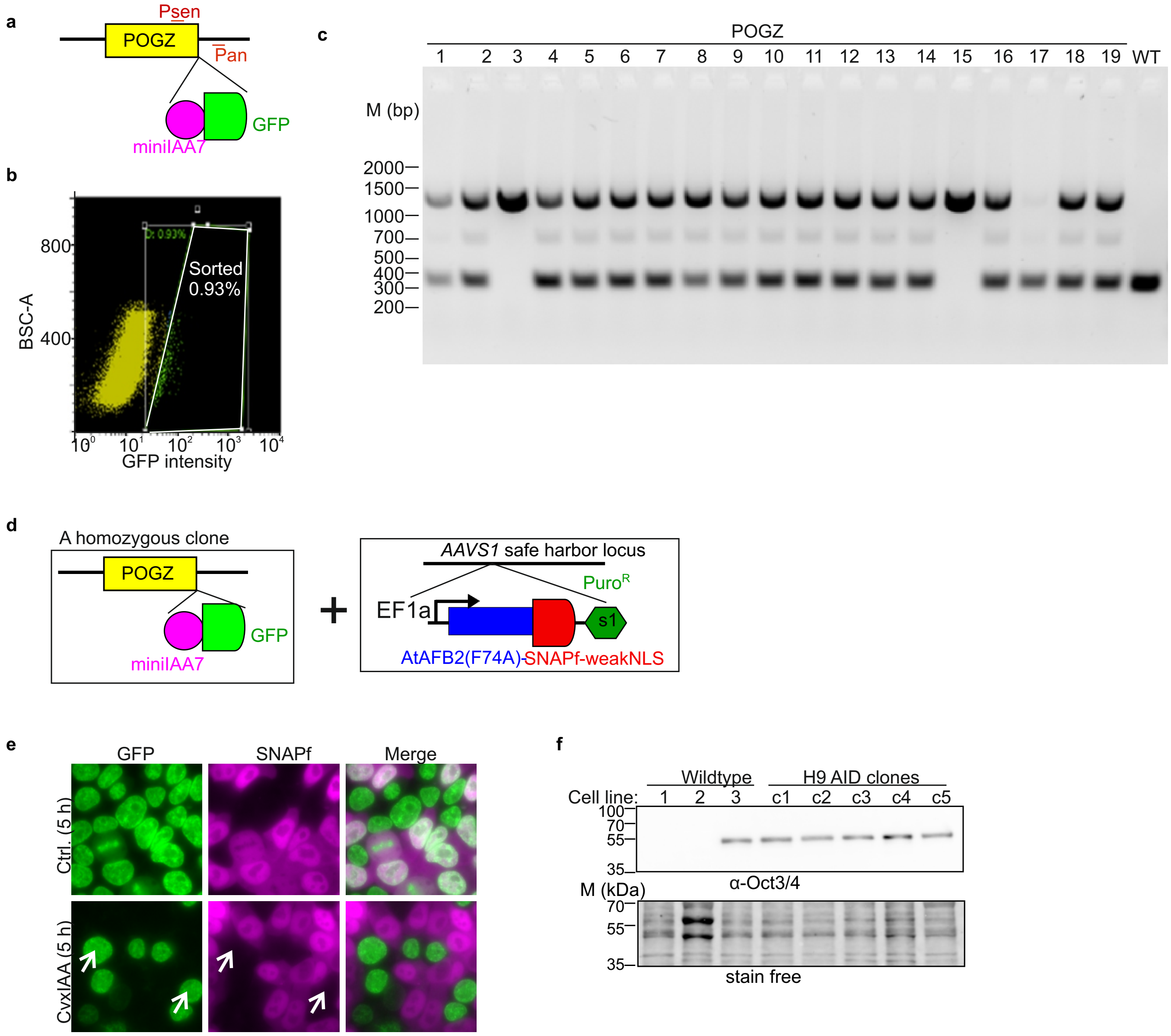


**Suppl. Fig. 8: generation of AID H9 cells with a conventional procedure.**

**a,** scheme showing endogenous POGZ tagging with miniIAA7-mEGFP at C-terminus. Pse and Pan for genomic PCR in (c).

**b,** FACS plot of analyzing POGZ tagging efficiency. Cells in the white frame (0.93%) were used for single-cell cloning.

**c,** genotyping PCR identifying POGZ-miniIAA7-mEGFP single-cell clones derived from sorted cells in (b).

**d,** scheme showing introduction of *At*AFB2(F74A)-SNAPf-weakNLS into a homozygous clone through *AAVS1* integration.

**e,** live cell images showing single-cell clones generated from (d) with or without 5 µM cvx_IAA induction for 5 h. Arrows indicate cells without *At*AFB2(F74A) expression and showing defective POGZ depletion after cvx_IAA induction. Representatives of 10 clones.

**f,** WB analysis of Oct3/4 expression in H9 AID clones. Wildtype 1-3: A431 (1), neuron differentiated from H9 (2) and H9 (3); c1-c5: H9 AID clones targeting NUP93 (c1), RANGAP1 (c2), SAC1 (c3), PEX3 (c4) and Seipin (c5). Representative of 2 AID clones for each target.
